# A postnatal molecular switch drives the activity-dependent maturation of parvalbumin interneurons

**DOI:** 10.1101/2024.04.08.588555

**Authors:** Monika Moissidis, Leyla Abbasova, Rafael Alis, Clémence Bernard, Yaiza Domínguez, Shenyue Qin, Audrey Kelly, Fazal Oozeer, Laura Mòdol, Fursham Hamid, Paul Lavender, Nuria Flames, Oscar Marín

## Abstract

Cortical neurons are specified during embryonic development but often only acquire their mature properties at relatively late stages of postnatal development. This delay in terminal differentiation is particularly prominent for fast-spiking parvalbumin-expressing (PV+) interneurons, which play critical roles in regulating the function of the cerebral cortex. We found that the maturation of PV+ interneurons is triggered by neuronal activity and mediated by the transcriptional cofactor peroxisome proliferator-activated receptor-gamma coactivator 1-alpha (PGC-1α). Developmental loss of PGC-1α prevents PV+ interneurons from acquiring unique structural, electrophysiological, synaptic, and metabolic features and disrupts their diversification into distinct subtypes. PGC-1α exerts its function as a master regulator of the differentiation of PV+ interneurons by directly controlling gene expression through a transcriptional complex that includes ERRγ and Mef2c. Our results uncover a molecular switch that translates neural activity into a specific transcriptional program promoting the maturation of PV+ interneurons at the appropriate developmental stage.

---

Understanding the molecular mechanisms controlling the diversification and terminal differentiation of neurons in the mammalian nervous system remains an immense challenge. This task is particularly daunting in the mammalian cerebral cortex, which contains a vast diversity of glutamatergic principal neurons and GABAergic interneurons^1–3^. Although cortical neurons are thought to be specified during embryonic development^4–7^, many do not express genes that regulate critical aspects of their function until relatively late stages of postnatal development. This observation suggests that the terminal differentiation of cortical neurons is linked to neuronal activity and the environmental signals the animals receive during postnatal development, as recently shown for certain types of excitatory cells^8^. However, since the maturation of cortical neurons is asynchronous, different cell types must have evolved specific molecular mechanisms translating environmental cues into signals for terminal differentiation at the appropriate developmental stage.

Fast-spiking interneurons expressing the calcium-binding protein parvalbumin (PV+) are critical regulators of cortical function^9,10^. They have unique electrophysiological characteristics associated with high metabolic demands, including fast action potential kinetics and ion conductances^10^. Most PV+ basket cells are also wrapped by perineuronal nets, a condensed form of the extracellular matrix that protects neurons from oxidative stress^11^ and regulates their plasticity^12^. Critically, PV+ interneurons make synapses on the somatic and perisomatic compartments of pyramidal cells, thereby exerting firm control over the firing of these cells^10^. The fast perisomatic inhibition of pyramidal cells by PV+ interneurons is required to generate gamma oscillations^13–15^, a high-frequency network rhythm essential for multiple cognitive operations often disrupted in neuropsychiatric disorders^16^.

Like other cells, PV+ interneurons acquire unique morphological, neurochemical, and electrophysiological features through the progressive selection of dedicated gene expression programs during development^4,5,17^. However, PV+ interneurons exhibit a particularly protracted period of maturation compared to other interneurons. In humans, for example, the expression of PV only increases dramatically after infancy, and the perineuronal nets covering these cells acquire their mature appearance around eight years of age^18,19^. Even in rodents, which exhibit a much more accelerated tempo of neuronal differentiation than primates^20,21^, PV+ interneurons only fully acquire the properties that enable their high-frequency action potentials well into the third week of postnatal life^22,23^. The maturation of PV+ interneurons critically regulates the transition of cortical activity from highly synchronous to sparse and decorrelated^24^, the closure of critical periods of heightened plasticity^25–27^, and the developmental emergence of complex cognitive abilities in mammals.

The maturation of PV+ interneurons occurs parallel to a rapid increase in glutamatergic synapses received by these cells^28,29^, suggesting that neuronal activity plays a role in this process. Consistent with this hypothesis, the density of glutamatergic synapses targeting PV+ interneurons in the adult cortex predicts cellular PV levels in both mice and humans^30,31^. Neuronal activity is also responsible for the secretion of brain-derived neurotrophic factor (BDNF) by excitatory neurons, which promotes PV+ interneuron development^32–34^. However, the precise molecular mechanisms through which neuronal activity may trigger this critical process in the development of cortical circuits remain to be elucidated.

Here, we investigated the cellular and molecular mechanisms regulating the terminal differentiation of PV+ interneurons in the mouse cerebral cortex. We found that the maturation of PV+ interneurons is modulated by neuronal activity and that this process requires the function of PGC-1α (peroxisome proliferator-activated receptor-gamma coactivator 1-alpha), a transcriptional cofactor upregulated by prospective PV+ interneurons postnatally. In other tissues, different isoforms of PGC-1α are involved in adaptive responses to change, a function that is thought to be mediated by their capacity to promote mitochondria biogenesis and energy metabolism^35,36^. In the brain, previous studies have shown that PGC-1α regulates the expression of PV in the adult cortex^37,38^, but its role in the protracted differentiation of PV+ interneurons, which is fundamental for circuit function, has not been explored. Our experiments demonstrate that PGC-1α is an activity-dependent master regulator of the maturation of PV+ cells, promoting the acquisition of adult features in these interneurons through transcriptional interactions involving estrogen-related receptor gamma (ERRγ) and myocyte-specific enhancer factor 2c (Mef2c).

## RESULTS

### PV+ interneuron maturation requires neuronal activity

To investigate whether the maturation of PV+ interneurons depends on neuronal activity, we increased the excitability of these cells immediately before the onset of PV expression using a chemogenetic approach based on Designer Receptors Exclusively Activated by Designer Drugs (DREADDs)^39^. In brief, we injected the primary somatosensory cortex (S1) of postnatal day (P) 0 *Lhx6-Cre* pups with a cocktail of Cre-dependent adeno-associated viruses (AAVs) expressing GFP and mutant G-protein-coupled receptors that induce neuronal activation (hM3Dq) following administration of the pharmacologically inert molecule clozapine-*N*-oxide (CNO). We titrated the hM3Dq virus to infect a relatively small fraction of interneurons in each animal (Figure S1A), thereby minimizing the possibility of broad network-induced changes (Figure S1B). We then injected CNO or vehicle twice daily from P10 to P12 and analyzed the impact of this manipulation at P12, focusing on layer (L) 2/3 (Figure 1A). We used *Lhx6-Cre* mice because cortical interneurons derived from the medial ganglionic eminence (MGE), including prospective PV+ and SST+ interneurons, express this transcription factor as soon as they become postmitotic^40^. Notably, SST is already expressed by migrating SST+ interneurons^41^, so prospective PV+ interneurons can be identified among the GFP+ (recombined) cells by the lack of SST expression (Figure 1B). We found that the intensity of PV and *Wisteria floribunda* Agglutinin (WFA), a marker of perineuronal nets, is higher in prospective PV+ interneurons expressing hM3Dq than in neighboring interneurons lacking hM3Dq expression (Figures 1B and 1C). We also verified that CNO alone does not influence the intensity of PV and WFA (Figure S1C).

**Figure 1.**
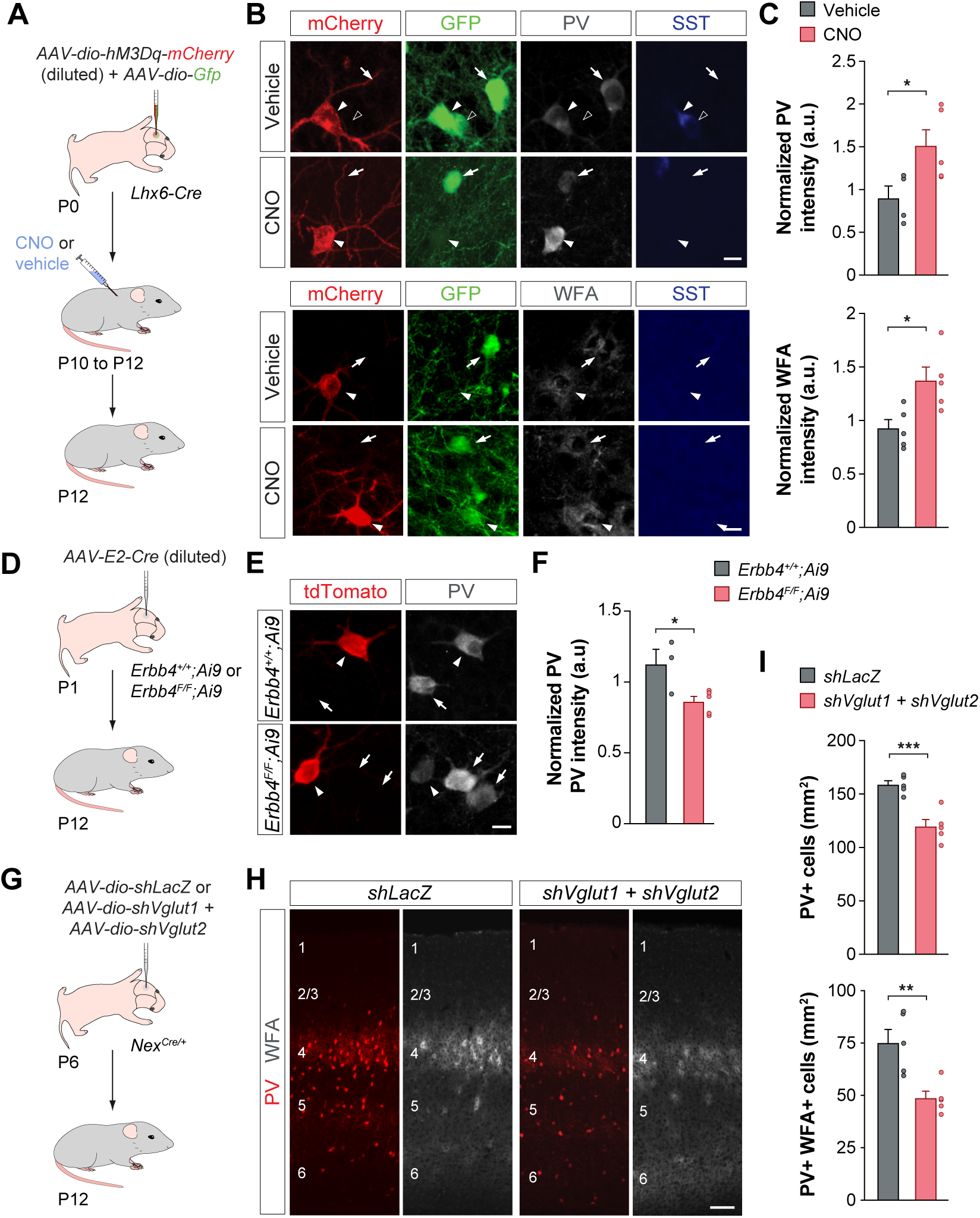
The maturation of cortical PV+ interneurons depends on neuronal activity. (A) Schematic representation of the experimental design. (B) PV and WFA expression in S1 L2/3 prospective PV+ cells (recognized by the absence of SST expression) expressing mCherry or GFP in control (vehicle) and experimental (CNO) conditions at P12. Arrows point at GFP+ PV+ cells, arrowheads point at mCherry+ PV+ cells, and open arrowheads point at GFP+ SST+ cell. (C) Quantification of PV and WFA levels in S1 L2/3 mCherry+ prospective PV+ cells normalized by PV and WFA levels in neighboring mCherry-GFP+ prospective PV+ cells at P12. Unpaired Student’s *t*-test, \**p* < 0.05. (D) Schematic representation of the experimental design. (E) PV expression in S1 L5/6 tdTomato+ PV+ cells in *ErbB4^+/+^;Ai9* and *ErbB4^F/F^;Ai9* mice at P12. Arrows and arrowheads point to uninfected and infected cells, respectively. (F) Quantification of PV levels in S1 L5/6 infected cells normalized by PV levels in neighboring uninfected cells at P12. Unpaired Student’s *t*-test, \**p* < 0.05. (G) Schematic representation of the experimental design. (H) PV and WFA expression in coronal sections through S1 from control (*shLacZ*) and experimental (*shVglut1 + shVglut2*) mice at P12. (I) Quantification of the density of PV+ and PV+ WFA+ cells in S1 from control (*shLacZ*) and experimental (*shVglut1 + shVglut2*) mice at P12. Unpaired Student’s *t*-test, \*\**p* < 0.01 and \*\*\**p* < 0.001. Data are presented as mean ± s.e.m. Scale bars, 10 µm (B and E) and 100 µm (H).

We wondered whether reducing the level of neural activity had the opposite effect on the maturation of PV+ interneurons. To test this hypothesis, we sparsely infected Lhx6+ interneurons by injecting S1 of P0 *Lhx6-Cre* pups with a cocktail of Cre-dependent AAVs expressing GFP and Cre-dependent AAVs encoding mutant G-protein-coupled receptors that reduce neuronal activation (hM4Di) following CNO administration. We then injected CNO or vehicle twice daily from P10 to P12 and analyzed the impact of this manipulation at P12 (Figure S1D). We focused our analysis on L5 because PV+ interneurons in this layer are at a more advanced stage of maturation than in L2/3 at this age. We found that the intensity of PV is decreased in prospective PV+ interneurons expressing hM4Di compared to neighboring infected cells lacking hM4Di expression (Figures S1E and S1F). Together, these experiments suggested that neuronal activity is necessary and sufficient to induce the maturation of PV+ interneurons.

Since pyramidal cells are the primary source of excitatory activity for cortical interneurons, we investigated whether the formation of glutamatergic synapses impacts the maturation of PV+ interneurons. To this end, we manipulated the density of glutamatergic synapses contacting prospective PV+ interneurons during early postnatal development. We and others have previously shown that the formation of these synapses requires the function of the tyrosine kinase receptor ErbB4^42–44^. To examine the effect of the loss of ErbB4 in a sparse population of prospective PV+ interneurons, we crossed *Erbb4^+/+^* and *Erbb4^F/F^*mice with *RCL^tdTomato/+^* (*Ai9*) reporter mice to label Cre-infected cells. We then injected S1 of P1 pups with AAVs expressing Cre recombinase under the control of the E2 enhancer, which directs Cre expression to putative PV+ cells^45^ (Figure 1D; see Methods for details). We titrated the virus to infect a relatively small fraction of interneurons in each animal (Figure S1G) and verified that, as expected, the loss of ErbB4 reduced the density of excitatory inputs contacting infected PV+ interneurons by P12 (Figure S1H). We found that tdTomato+ PV+ interneurons in *Erbb4^F/F^;Ai9* mice had lower levels of PV intensity than neighboring uninfected PV+ cells, a phenotype that was not observed in *Erbb4^+/+^;Ai9* mice (Figure 1E and 1F). These experiments reinforced that neuronal activity, mediated by glutamatergic synapses, is critical for the maturation of PV+ interneurons.

To strengthen this idea, we reduced glutamatergic neurotransmission globally in the neocortex by downregulating the expression of the glutamate transporters Vglut1 and Vglut2 in pyramidal cells. To this end, we designed conditional short-hairpin RNA (shRNA) vectors against *Slc17a6* and *Slc17a7*, which encode Vglut1 and Vglut2, respectively. We then generated Cre-dependent AAVs expressing the shRNA constructs and injected them into S1 of P6 *Nex^Cre/+^* pups, in which Cre is expressed by all principal cells (Figure 1G). We confirmed the ability of *shVglut1* and *shVglut2* to downregulate the expression of the corresponding target genes using mice injected with Cre-dependent AAVs expressing *shLacZ* as controls (Figures S1I and S1J). Because glutamatergic neurotransmission is required for interneurons to survive in the early postnatal cortex^46–48^, we timed these experiments to avoid the period of programmed cell death in interneurons. We verified that the downregulation of Vglut1 and Vglut2 did not impact the density of Lhx6+ cells at P12 (Figure S1K). In contrast, we found that the density of cells expressing detectable levels of PV and perineuronal nets was decreased in mice with reduced glutamatergic neurotransmission compared to controls (Figures 1H and 1I). Altogether, our experiments indicated that neuronal activity is critical for the maturation of PV+ interneurons.

### PGC-1α mediates the effect of activity in the maturation of PV+ interneurons

We next explored the molecular mechanism through which neuronal activity may influence the maturation of PV+ interneurons. Previous work has shown that PGC-1⍺, a transcriptional cofactor linking different environmental stimuli to adaptive responses in multiple tissues, is required for PV expression in the adult cortex^37^. We investigated whether the expression of *Ppargc1a*, the gene encoding PGC-1⍺, precedes the onset of PV expression in the postnatal cortex. We found that *Ppargc1a* begins to be expressed in the neocortex around P5, and from P7 onwards – before the onset of PV expression and perineuronal net formation – it concentrates in sparse somas spanning L2 to L6 (Figures 2A and 2B). We observed that most cortical cells containing high *Ppargc1a* mRNA levels are MGE-derived interneurons (Figures S2A and S2B), and the proportion of MGE-derived interneurons expressing *Ppargc1a* increases progressively until reaching a plateau at P12 (Figure S2C). We also analyzed the proportion of MGE-derived cells expressing PGC-1⍺ during the onset of PV expression in the neocortex. Most MGE-derived cells expressing PGC-1⍺ are prospective PV+ interneurons (GFP+ SST-cells), and their proportion increases between P10 and P12 (Figures 2C and 2D). Interestingly, PGC-1⍺ also concentrates in the nucleus of prospective PV+ interneurons during this period (Figures 2E and 2F). In contrast, the proportion of SST+ cells expressing PGC-1⍺ is relatively low and remains constant during this period (Figures 2C and 2D). These observations revealed that PGC-1⍺ expression in the neocortex concentrates in prospective PV+ interneurons before the onset of PV expression.

**Figure 2.**
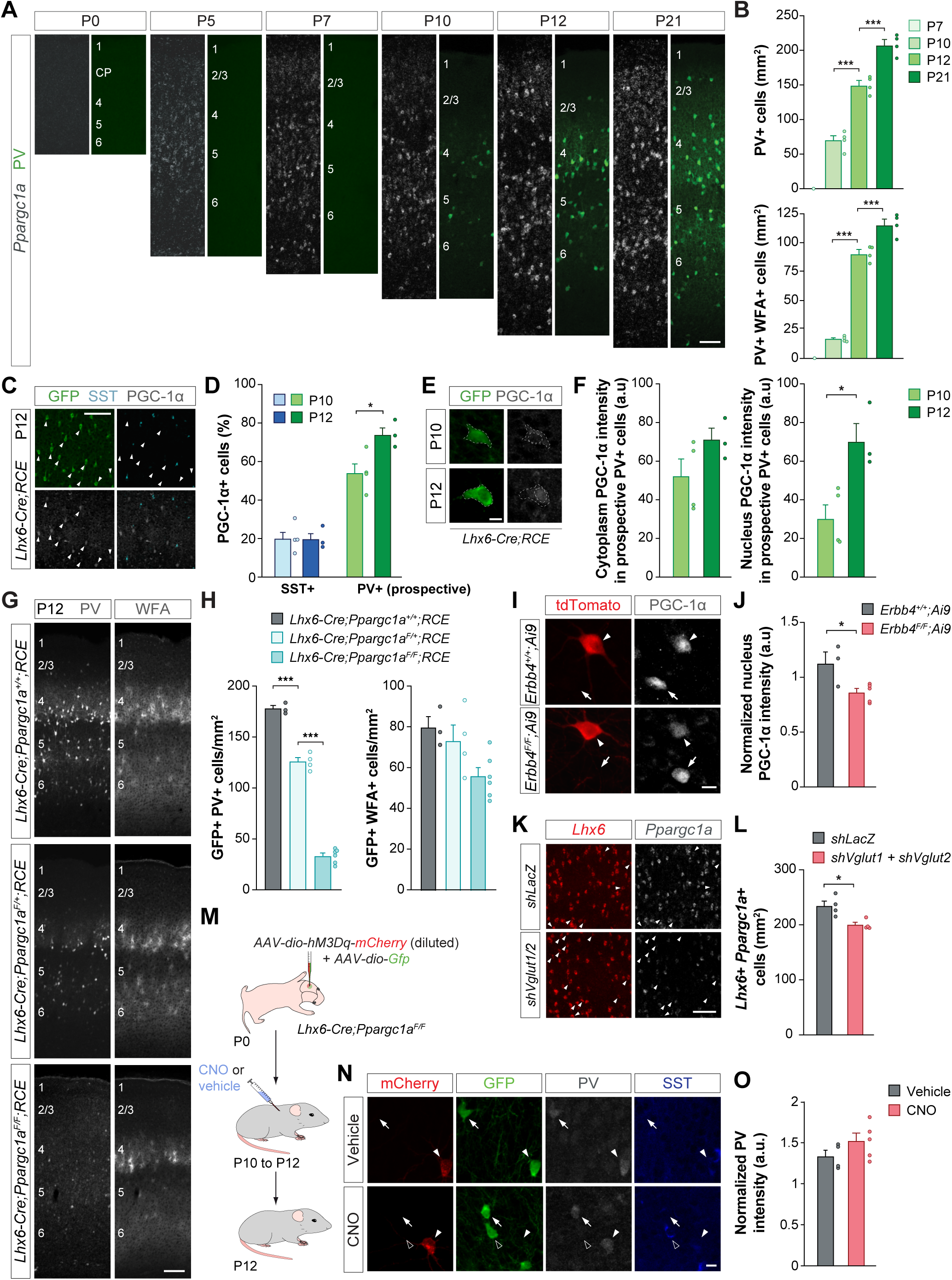
The maturation of cortical PV+ interneurons depends on the activity-dependent expression of PGC-1⍺. (A) PV and *Ppargc1a* expression in coronal sections through S1 of wild-type mice at P0, P5, P7, P10, P12 and P21. (B) Quantification of the density of PV+ and PV+ WFA+ cells in S1 during postnatal development. One-way ANOVA, \*\*\**p* < 0.001. (C) PGC-1⍺ expression in SST+ cells and prospective PV+ interneurons in S1 of *Lhx6-Cre;RCE* mice at P12. Arrowheads point at prospective PV+ interneurons expressing PGC-1⍺. (D) Quantification of the proportion of SST+ and prospective PV+ interneurons expressing PGC-1⍺ at P10 and P12 in S1 of *Lhx6-Cre;RCE* mice. Two-way ANOVA followed by Tukey’s multiple correction tests, \**p* < 0.05. (E) PGC-1⍺ expression in the cytoplasm and nucleus of MGE-derived cells in *Lhx6-Cre;RCE* mice at P10 and P12. (F) Quantification of PGC-1⍺ levels in the cytoplasm and nucleus of prospective PV+ interneurons at P10 and P12 in S1 of *Lhx6-Cre;RCE* mice. Unpaired Student’s *t*-test, \**p* < 0.05. (G) PV and WFA expression in coronal sections through S1 from control, conditional heterozygous, and conditional homozygous *Ppargc1a* mutant mice at P12. (H) Quantification of the density of GFP+ PV+ cells and GFP+ WFA+ cells in S1 of control, conditional heterozygous, and conditional homozygous *Ppargc1a* mice at P12. One-way ANOVA followed by Tukey’s multiple correction tests, \*\*\**p* < 0.001. (I) PGC-1⍺ expression in infected PV+ cells from *ErbB4^+/+^;Ai9* and *ErbB4^F/F^;Ai9* mice at P12. Arrows and arrowheads point at uninfected and infected cells, respectively. (J) Quantification of nuclear PGC-1⍺ levels in infected PV+ cells normalized by nuclear PGC-1⍺ levels in neighboring uninfected PV+ cells. Unpaired Student’s *t*-test, \**p* < 0.05. (K) *Ppargc1a* expression in *Lhx6+* cells from control (*shLacZ*) and experimental (*shVglut1 + shVglut2*) mice at P12. Arrowheads point at *Lhx6+ Ppargc1a*-cells. (L) Quantification of the density of *Lhx6+ Ppargc1a+* cells in S1 from control (*shLacZ*) and experimental (*shVglut1 + shVglut2*) mice at P12. Unpaired Student’s *t*-test, \**p* < 0.05. (M) Schematic representation of the experimental design. (N) PV expression in prospective PV+ cells (recognized by the absence of SST expression) expressing mCherry or GFP in control (vehicle) and experimental (CNO) conditions at P12. Arrows point at GFP+ PV+ cells, arrowheads point at mCherry+ PV+ cells, and open arrowheads point at GFP+ SST+ cell. (O) Quantification of PV levels in S1 L2/3 mCherry+ prospective PV+ cells normalized by PV levels in neighboring mCherry-GFP+ prospective PV+ cells at P12. Unpaired Student’s *t*-test, not significant. Data are presented as mean ± s.e.m. Scale bars, 10 µm (E, I and N), 100 µm (A, C, K and G).

To investigate whether PGC-1⍺ is necessary for the maturation of PV+ interneurons, we generated conditional mutants in which we deleted *Ppargc1a* specifically from MGE-derived cells. To this end, we bred *Lhx6-Cre* mice with mice carrying floxed *Ppargc1a* alleles and with *RCL^Gfp/+^* (RCE) reporter mice to label Cre-expressing cells and verified the effective loss of PGC-1⍺ in *Lhx6-Cre;Ppargc1a^F/F^;RCE* mice by P12 (Figure S2D and S2G). We found that the loss of PGC-1⍺ does not impact the density or the laminar distribution of MGE-derived interneurons (Figures S2E and S2F). However, mice lacking one or two *Ppargc1a* alleles from MGE-derived interneurons exhibit a dose-dependent loss of PV expression, which is nearly absent in conditional homozygous *Ppargc1a* mutant mice (Figures 2G, 2H, and S2H). Conditional mutant mice also have fewer perineuronal nets surrounding PV+ cells than controls, specifically in L4, where perineuronal nets are usually most abundant at P12 (Figures 2G, 2H, and S2H). Importantly, we found no difference in the density of SST+ interneurons in conditional homozygous *Ppargc1a* mutant mice compared to controls (Figures S2J and S2K), suggesting that the loss of PGC-1⍺ does not affect the maturation of these cells and does not change the fate of PV+ interneurons into an SST+ interneuron fate.

We assessed whether PGC-1⍺ mediates the role of activity in the maturation of PV+ interneurons. To this end, we first examined whether reducing the density of excitatory inputs contacting infected PV+ interneurons (Figure 1D) or reducing glutamatergic neurotransmission globally in the neocortex (Figure 1G) impacts the expression of PGC-1⍺ in prospective PV+ cells at P12. We found that both manipulations, which decrease the levels of PV and WFA in prospective PV+ cells (Figures 1E, 1F, 1H, and 1I), diminish nuclear PGC-1⍺ levels in these cells and the number of *Lhx6*+ cells expressing *Ppargc1a* (Figures 2I–2L). Next, we performed a new series of experiments in which we increased the activity of prospective PV+ cells between P10 and P12 using activating DREADDs, but, on this occasion, we carried out the manipulation in *Lhx6-Cre;Ppargc1a^F/F^* mice (Figure 2M). In contrast to the results in wild-type mice (Figures 1B and 1C), we found no differences in the expression of PV and WFA between vehicle- and CNO-injected mutant mice (Figures 2N, 2O, and S2L). These experiments revealed that PGC-1⍺ mediates the impact of neuronal activity on the maturation of PV+ interneurons during early postnatal development.

### PGC-1α controls the terminal differentiation of PV+ interneurons

Our previous experiments suggested that PGC-1⍺ plays a critical role in the maturation of PV+ interneurons during early postnatal development. However, although PV and WFA are essential markers of mature PV+ interneurons^10^, it is unclear whether this factor regulates a broader molecular program required for the terminal differentiation of cortical PV+ interneurons. To explore this possibility, we analyzed gene expression in MGE-derived interneurons from control (*Lhx6-Cre;Ppargc1a^+/+^*) and conditional *Ppargc1a* mutants (*Lhx6-Cre;Ppargc1a^F/F^*) using single-cell RNA sequencing experiments (scRNA-seq). The maturation of PV+ interneurons continues into the fourth week of postnatal development in S1, with the slow acquisition of mature electrophysiological properties^28^. We thus performed these experiments at P21 to identify gene differences that might not be detected at earlier stages. To this end, we injected Cre-dependent AAVs expressing tdTomato under the control of the neuron-specific human synapsin 1 gene promoter^49^ into S1 of P1 pups. We then generated single-cell suspensions from S1 at P21, isolated the labeled cells using fluorescence-activated cell sorting, and performed scRNA-seq (Figure 3A). We took this approach to avoid isolating endothelial cells, which are also recombined in *Lhx6-Cre* mice^50^ (Figures S2D and S2E).

**Figure 3.**
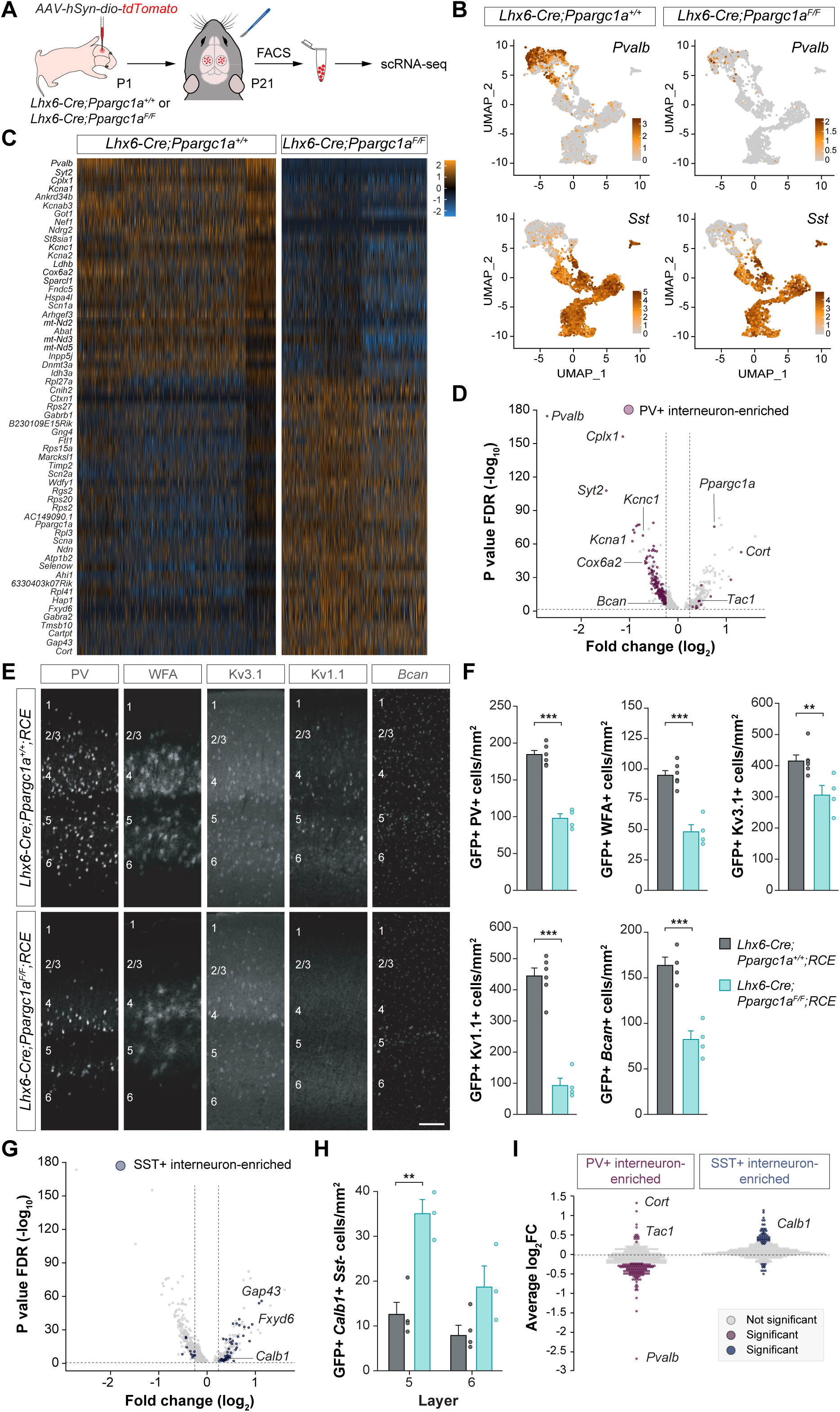
PGC-1⍺ is a master regulator of the maturation of cortical PV+ interneurons. (A) Schematic representation of the experimental design. (B) UMAP plots illustrating the expression of *Pvalb* and *Sst* in control and conditional homozygous *Ppargc1a* mice at P21. (C) Heatmap representing differential gene expression between PV+ interneurons in control and conditional homozygous *Ppargc1a* mice at P21. (D) Volcano plot illustrating differentially expressed genes (DEGs) between PV+ interneurons in control and conditional homozygous *Ppargc1a* mice at P21. A threshold at log_2_FC > ±0.25 was set. Genes normally enriched in PV+ interneurons are highlighted in magenta. (E) PV, WFA, Kv3.1, Kv1.1, and *Bcan* expression in coronal sections through S1 of control and conditional homozygous *Ppargc1a* mice at P21. (F) Quantification of the density of GFP+ PV+, GFP+ WFA+, GFP+ Kv3.1+, GFP+ Kv1.1+, and GFP+ *Bcan*+ cells in S1 of control and conditional homozygous *Ppargc1a* mice at P21. Unpaired Student’s *t*-test, \**p* < 0.05, \*\*\**p* < 0.001. (G) Volcano plot illustrating DEGs between PV+ interneurons in control and conditional homozygous *Ppargc1a* mice at P21. A threshold at log_2_FC > ±0.25 was set. Genes normally enriched in SST+ interneurons are highlighted in blue. (H) Quantification of the density of GFP+ *Calb1*+ *Sst*-cells in L5 and L6 of S1 in control and conditional homozygous *Ppargc1a* mice at P21. Two-way ANOVA followed by Tukey’s multiple correction tests, \*\**p* < 0.01. (I) Quantification of expression changes (average log_2_FC) in PV+ interneurons between control and conditional homozygous *Ppargc1a* mice at P21 for genes normally enriched in PV+ interneurons or SST+ interneurons. A threshold at log_2_FC > ±0.25 was set. Significant changes in expression are highlighted in magenta and blue for genes normally enriched in PV+ and SST+ interneurons, respectively. Data are presented as mean ± s.e.m. Scale bar, 100 µm (E).

Following quality control (Figure S3), we isolated 5,268 control and 3,139 mutant MGE-derived interneurons from three highly consistent experimental replicas for each genotype (Figures S4A and S4B). We then merged the replicates, integrated control and mutant conditions, and classified the cells into PV+ and SST+ interneurons based on *Pvalb* and *Sst* gene expression (Figures 3B and S4C). We found that PV+ and SST+ interneurons lacking *Ppargc1a* cluster with their corresponding control cells (Figure 3B), suggesting that PGC-1⍺ does not control the fate of MGE-derived interneurons.

We performed differential gene expression (DGE) analysis between control and conditional mutant PV+ interneurons and identified genes modified by the absence of *Ppargc1a* in these cells. We found that many highly characteristic genes of PV+ interneurons were downregulated in mutant cells compared to control neurons (Figure 3C). Because PV+ and SST+ have a common developmental origin and shared transcriptional programs^5^, we identified PV-enriched genes by comparing the transcriptome of control PV+ and SST+ interneurons at P21 (Figure S4D) and analyzed their expression in mutant PV+ cells. We found that many PV-enriched genes were downregulated in PV+ cells lacking PGC-1⍺, including *Pvalb*, *Cplx1*, *Syt2*, *Kcnc1, Kcna1*, and *Bcan*, and we verified these results by immunohistochemistry and in situ hybridization (Figures 3D–3F). These genes encode proteins controlling a broad range of critical properties in mature PV+ interneurons, comprising neurochemical (e.g., PV and brevican), electrophysiological (e.g., potassium voltage-gated channel, Shaw-related subfamily, member 1 [Kv3.1] and potassium voltage-gated channel, shaker-related subfamily, member 1 [Kv.1.1]), and synaptic (e.g., complexin 1 and synaptotagmin 2) features. We noticed that comparatively fewer genes were upregulated in mutant PV+ interneurons compared to controls (Figure 3D). We found *Ppargc1a* among the upregulated genes (transcripts lacking the floxed exons 3-5), suggesting compensatory effects for the lack of PGC-1⍺.

Neuronal maturation does not only require the implementation of specific effector gene programs. In addition, increasing evidence supports the importance of active repression to avoid the expression of alternative fates of related developmental lineages^51^. We wondered whether PV+ interneurons lacking PGC-1⍺ express genes normally enriched in SST+ interneurons, with whom they share a common developmental lineage. To this end, we first identified SST-enriched genes by comparing the transcriptome of control PV+ and SST+ interneurons at P21 (Figure S4D) and analyzed their expression in mutant PV+ cells. We found that several SST-enriched genes were upregulated in PV+ cells lacking PGC-1⍺. These include the gene encoding calbindin (*Calb1*), a calcium-binding protein frequently found in deep-layer SST+ interneurons^52^, and we verified this result by in situ hybridization (Figures 3G and 3H). Looking at gene expression changes more globally, we observed that PV-enriched genes were more frequently downregulated, and SST-enriched genes were more frequently upregulated in PV+ interneurons lacking PGC-1⍺ compared to control cells (Figure 3I). To investigate whether this pattern also applies to other cortical interneurons from more distant developmental lineages, we used a published dataset^53^ to identify genes enriched in those cells and examined their expression in PV+ interneurons lacking PGC-1⍺. Using this alternative approach, we verified that PV-enriched genes are significantly downregulated in the absence of PGC-1⍺. We found that some genes normally enriched in other interneuron subclasses are differentially expressed in mutant PV+ interneurons, but the expression levels of the overall populations of subclass-enriched genes are only significantly upregulated for SST-enriched genes (Figure S4E).

PV+ interneurons have considerable metabolic demands due to their fast-spiking characteristics, and they contain considerably higher numbers of mitochondria than other interneurons and pyramidal cells^54–56^. Since PGC-1⍺ is a master regulator of mitochondrial biogenesis in other cellular contexts^57–59^, we wondered whether the loss of *Ppargc1a* may particularly disrupt the metabolism of PV+ interneurons. Pathway enrichment analysis comparing genes differentially expressed in PV+ and SST+ interneurons revealed that terms associated with metabolic pathways are specifically affected in mutant PV+ interneurons but not in mutant SST+ cells (Figure 4A). Several genes associated with mitochondrial function are normally enriched in PV+ interneurons and are downregulated in conditional *Ppargc1a* mutants, such as the gene encoding the cytochrome c oxidase, subunit 6a2, *Cox6a2* (Figures 3D, 4B, and 4C). We thus examined whether the loss of PGC-1⍺ affects the number of mitochondria in MGE-derived interneurons. To this end, we injected S1 of P1 control or *Ppargc1a* conditional mutants with AAVs expressing a protein to tag mitochondria, mito-EmGFP^60^, and analyzed their abundance in PV+ and SST+ cells at P21 (Figure 4D). We confirmed that control PV+ interneurons already have more mitochondria than SST+ cells at P21 (Figures 4E and 4F). Remarkably, PGC-1⍺ seems to be directly involved in generating this surplus of mitochondria in PV+ interneurons because the loss of *Ppargc1a* specifically affects the number of mitochondria in these cells but not in SST+ interneurons (Figures 4E and 4G).

**Figure 4.**
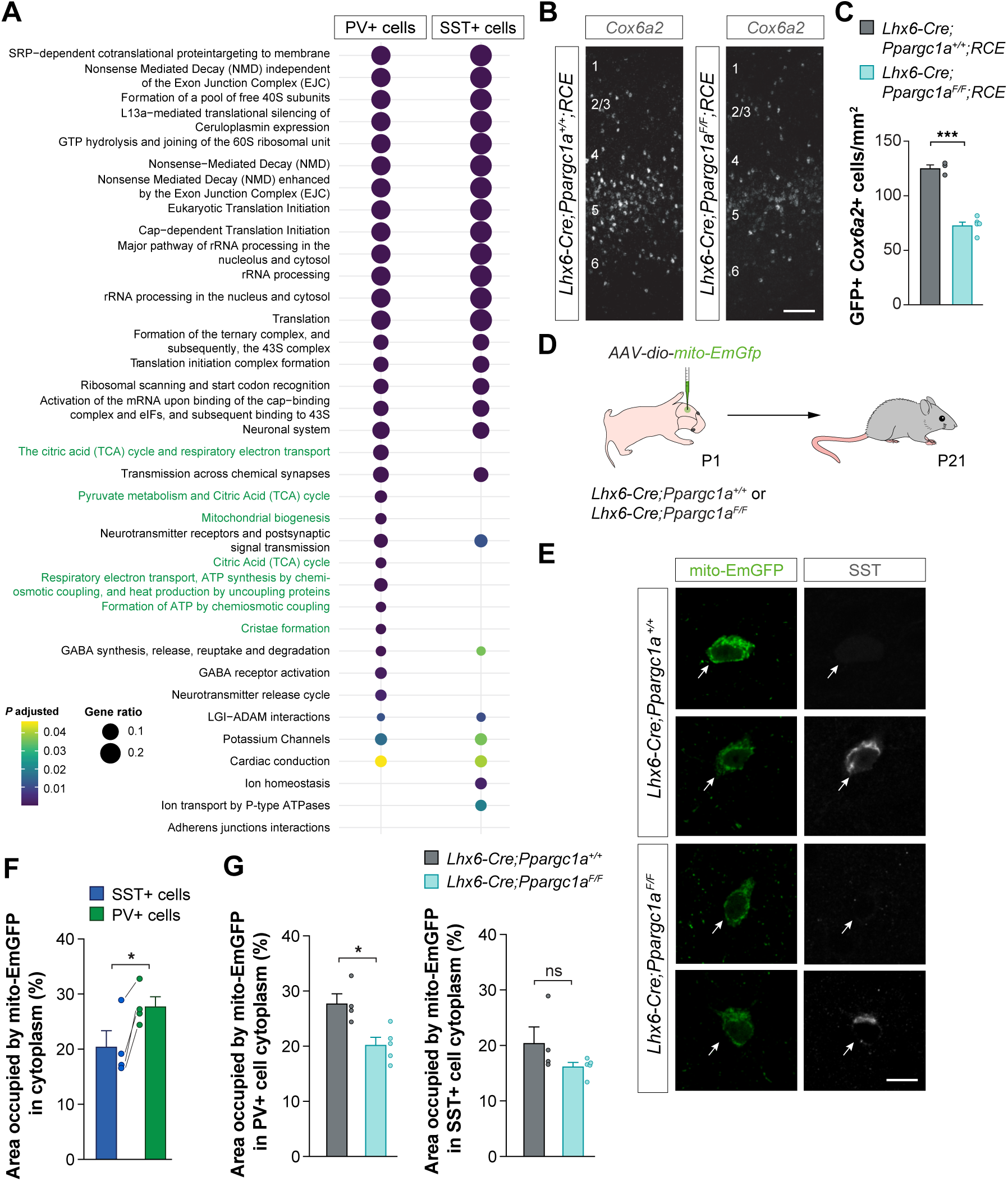
PGC-1⍺ regulates mitochondrial function in cortical PV+ interneurons. (A) Pathway enrichment analysis plot depicting the different cellular pathways affected by the absence of PGC-1⍺ in conditional homozygous PV+ and SST+ interneurons compared to controls. Pathways associated with mitochondrial function are highlighted in green. (B) *Cox6a2* expression in coronal sections through S1 of control and conditional homozygous *Ppargc1a* mice at P21. (C) Quantification of the density of GFP+ *Cox6a2*+ cells in S1 of control and conditional homozygous *Ppargc1a* mice at P21. Unpaired Student’s *t*-test, \*\*\**p* < 0.001. (D) Schematic representation of the experimental design. (E) mito-EmGFP expression in the soma of MGE-derived cells in S1 of control and conditional homozygous *Ppargc1a* mice at P21. PV+ interneurons are recognized by the absence of SST expression. (F) Quantification of the area occupied by mito-EmGFP signal compared to the total cytoplasmic area in S1 L2-4 PV+ and SST+ cortical interneurons at P21. Paired Student’s *t*-test, \**p* < 0.05. (G) Quantification of the area occupied by mito-EmGFP signal compared to the total cytoplasmic area in S1 L2-4 PV+ and SST+ cortical interneurons of control and conditional homozygous *Ppargc1a* mice at P21. Unpaired Student’s *t*-test, \**p* < 0.05. Data are presented as mean ± s.e.m. Scale bars, 10 µm (E), 100 µm (B).

Altogether, the results of the scRNA-seq experiments suggested that PGC-1⍺ is required for the expression of an extensive molecular program that controls the terminal differentiation of PV+ interneurons. In addition to promoting the expression of many critical factors in these cells, this molecular program may involve repressing genes normally found in SST+ interneurons.

### PGC-1α directly regulates the expression of target genes in PV+ interneurons

PGC-1⍺ may control the expression of target genes in PV+ interneurons by directly interacting at their promoters and enhancer sites or by modulating gene expression indirectly. For example, recent work suggests that mitochondria metabolism sets the tempo of neuronal maturation^60^, so disruption of mitochondria biogenesis in PV+ interneurons lacking PGC-1⍺ could perhaps be responsible for slowing down the differentiation of these cells. To test this hypothesis, we first examined whether the maturation of PV+ interneurons is simply delayed or permanently disrupted by analyzing the expression of PV and WFA in control and conditional *Ppargc1a* mutants at P60. We found that loss of PGC-1⍺ permanently impairs the maturation of PV+ cells across the cortex except for interneurons in L4, which have reached comparable levels to control mice by P60 (Figure S5). These observations revealed that the maturation of many PV+ interneurons is not simply delayed in the absence of PGC-1⍺, although some PV+ cells seem to recruit alternative programs to circumvent the loss of *Ppargc1a*.

The previous experiments suggested that PGC-1⍺ actively controls the expression of genes required for the maturation of PV+ interneurons. To determine whether PGC-1⍺ directly regulates the expression of these genes, we performed CUT&RUN experiments. In brief, we dissected S1 from wild-type mice at P12, prepared nuclear suspensions, and conducted CUT&RUN assays using an antibody against PGC-1⍺ (Figure 5A). We found that PGC-1⍺ binds at the promoter of genes or at distal intergenic and intronic regions that may correspond to enhancer sites (Figure S6A). Remarkably, we observed that 75% of the genes whose expression is altered in mutant PV+ cells are directly bound by PGC-1⍺ in cortical tissue, including PV- and SST-enriched markers and many metabolic genes (Figure 5B). For instance, PGC-1⍺ binds the promoters of *Bcan*, *Cplx1*, and *Kcna1* (Figure 5C), genes highly enriched in PV+ interneurons and essential for acquiring their unique properties. PGC-1⍺ also binds at enhancer sites for *Kcnc1* and at enhancer sites^61^ potentially associated with *Pvalb* and *Cox6a2*, genes highly specific to PV+ interneurons (Figure 5C). In addition, PGC-1⍺ seems to be able to modulate its own expression in the neocortex by binding the *Ppargc1a* brain-specific promoter^62^ (Figure 5C). Remarkably, PGC-1⍺ binds a large fraction of genes encoding metabolic (e.g., *Cox8a*, *Ldhb*, *Atp1b2*) and ribosomal (e.g., *Rpl30*, *Rpl41*, *Rps5, Rps26*) proteins whose expression is dysregulated in PV+ and SST+ interneurons lacking *Ppargc1a* (Figures 4A, S6B, and S6C).

**Figure 5.**
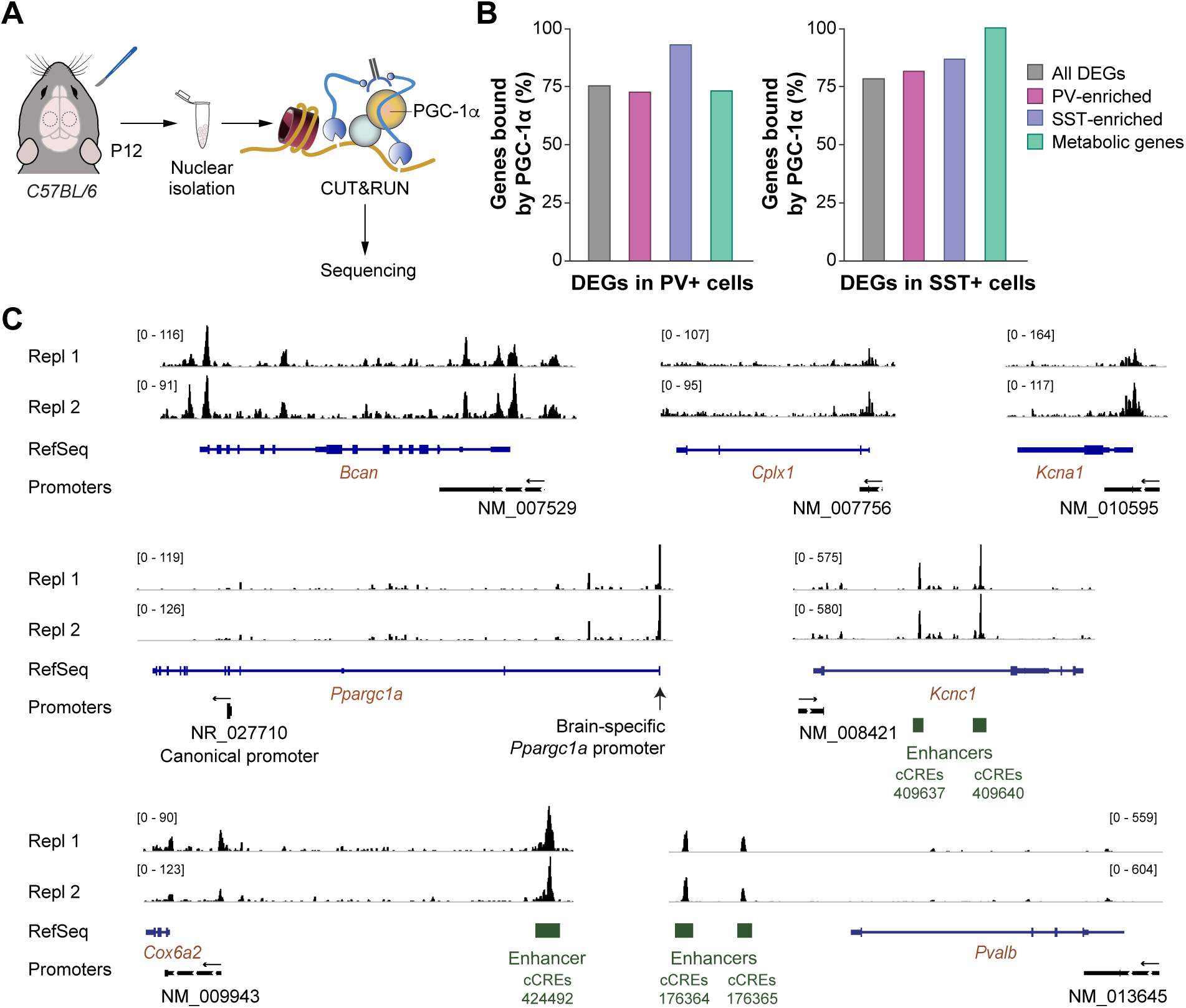
PGC-1⍺ regulates the maturation of cortical PV+ interneurons through direct transcriptomic regulation. (A) Schematic representation of the experimental design. (B) Plots depicting the proportion of DEGs between control and conditional homozygous *Ppargc1a* PV+ cells and SST+ cells, directly bound at their locus by PGC-1⍺. All DEGs (grey), PV-enriched genes (pink), SST-enriched genes (purple), and genes associated with metabolic function (green) were assessed. (C) Genome browser views of the genomic loci of the PV-enriched genes *Bcan, Cplx1, Kcna1, Ppargc1a, Kcnc1, Cox6a2,* and *Pvalb*. The black peaks represent the read coverage following PGC-1⍺ CUT&RUN in two replicates at P12. Gene promoters and enhancers, and the size scale of the peaks, are annotated. The cis regulatory elements (CRE) were identified using the database from Li and colleagues^61^. The arrows above the promoters indicate the direction of transcription.

PGC-1⍺ does not directly bind DNA and modulates gene expression by interacting with transcription factors and chromatin modifiers^36,63^. For instance, members of the estrogen-related receptor (ERR) family are known coactivators of PGC-1⍺ in several tissues^64^. Among them, ERR⍺ has been suggested to cooperate with PGC-1⍺ to regulate *Pvalb* expression in the adult cortex^65^. At P12 and P21, however, *Esrrg* mRNA is more abundantly expressed in the neocortex and is more specifically expressed in PV+ interneurons than *Esrra* (Figures S7A, S7B, and S7E), and it extensively colocalizes with *Ppargc1a* (Figures S7C and S7F). We also observed that 75% of PV+ interneurons express ERRγ while only 10% of SST+ interneurons do at P21 (Figures S7D and S7G). This specificity of ERRγ expression seems to be directly controlled by PGC-1⍺ as it binds the *Esrrg* locus in cortical cells at P12, and ERRγ levels are dramatically decreased in conditional *Ppargc1a* mutants (Figures 6A–6C).

**Figure 6.**
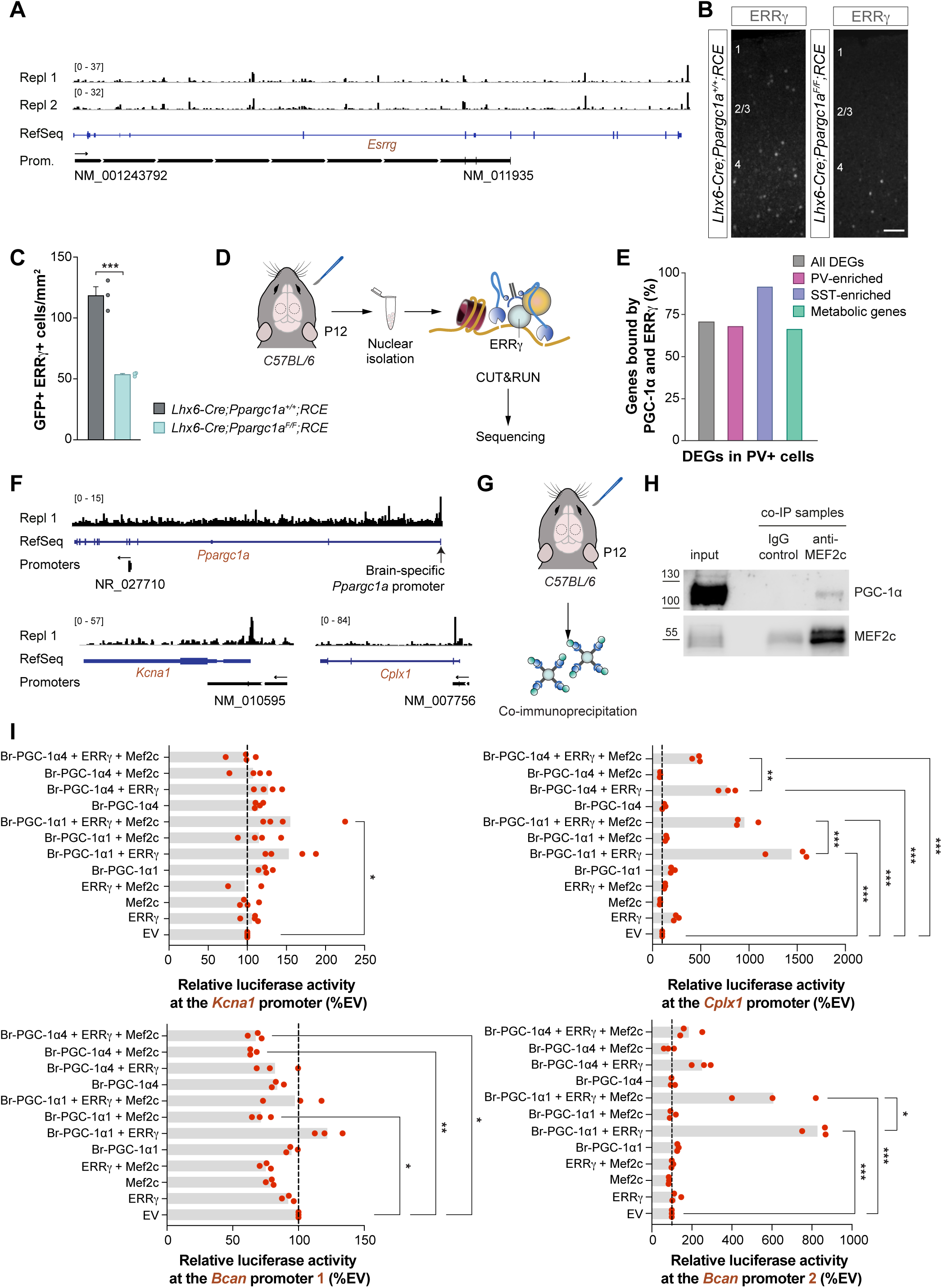
PGC-1⍺, ERRγ and Mef2C directly regulate gene expression in PV+ interneurons. (A) Genome browser view of the *Esrrg* genomic locus. The black peaks represent the read coverage following PGC-1⍺ CUT&RUN in two replicates at P12. Gene promoters and enhancers, and the size scale of the peaks, are annotated. The arrows above the promoters indicates the direction of transcription. (B) ERRγ expression in coronal sections through S1 of control and conditional homozygous *Ppargc1a* mice at P21. (C) Quantification of the density of GFP+ ERRγ+ cells in S1 of control and conditional homozygous *Ppargc1a* mice at P21. Unpaired Student’s *t*-test, \*\*\**p* < 0.001. (D) Schematic representation of the experimental design. (E) Plots depicting the proportion of differentially expressed genes (DEGs) between control and conditional homozygous *Ppargc1a* PV+ cells, directly bound at their locus by PGC-1⍺ and ERRγ. All DEGs (grey), PV-enriched genes (pink), SST-enriched genes (purple), and genes associated to metabolic function (green) were assessed. (F) Genome browser view of the genomic locus of *Ppargc1a, Kcna1,* and *Cplx1*. The black peaks represent the read coverage following ERRγ CUT&RUN at P12. Gene promoters and the size scale of the peaks, are annotated. The arrows above the promoters indicate the direction of transcription. (G) Schematic representation of the experimental design. (H) Co-immunoprecipitation of Mef2c identifies the long-isoform of PGC-1⍺ as a binding partner. (I) Luciferase assay experiments using selected DNA sequences of *Kcna1*, *Cplx1* and *Bcan* promoters bound by PGC-1⍺. Luciferase activity was assessed in the presence of different combinations of Mef2c, ERRγ, and the brain-specific short (PGC-1⍺4) and long (PGC-1⍺1) isoforms of PGC-1⍺. Data are presented as mean ± s.e.m. Scale bar, 100 µm (B).

These previous results suggested that ERRγ is a more likely coactivator of PGC-1⍺ than ERR⍺ to regulate the terminal differentiation of PV+ interneurons and induce the expression of PV-enriched genes in the early postnatal neocortex. To test this hypothesis, we dissected S1 from wild-type mice at P12, prepared nuclear suspensions, and conducted CUT&RUN assays using an antibody against ERRγ (Figure 6D). We found that approximately half of the loci (52.3%) bound by PGC-1⍺ in cortical neurons are also bound by ERRγ. Like PGC-1⍺, ERRγ binds at the promoter of genes or at distal intergenic and intronic regions that may correspond to enhancer sites (Figure S7H). We observed that 75% of the genes whose expression is altered in mutant PV+ interneurons are directly bound by ERRγ in cortical tissue, and most of these genes are also bound by PGC-1⍺ (Figure 6E). For example, ERRγ binds the promoters of *Kcna1*, *Cplx1*, and the brain-specific promoter of *Ppargc1a* (Figure 6F). These experiments revealed that PGC-1⍺ and ERRγ cooperate in regulating the terminal differentiation of PV+ interneurons in the early postnatal neocortex.

We performed a motif enrichment analysis at PGC-1⍺ binding loci to identify additional factors that may contribute to regulating PV-enriched genes. This analysis revealed that members of the myocyte enhancer factor 2 (Mef2) family are potential partners of PGC-1⍺ in the neocortex (Figure S6D). Importantly, Mef2c has been previously involved in the differentiation of PV+ interneurons^66,67^ and is expressed by most MGE-derived interneurons at P12 (Figures S8A–S8C). We found that Mef2c interacts with the long isoform of PGC-1⍺ (PGC-1⍺1) at P12 (Figure 6H), even though the short isoform (PGC-1⍺4) is much more abundant in the neocortex at this age (Figure S8D). This result revealed that Mef2c is another potential cofactor of PGC-1⍺ in PV+ interneurons.

Since Mef2c is already expressed in cortical neurons at embryonic stages^66,67^, we wondered whether this transcription factor may regulate the expression of PGC-1⍺ in developing PV+ interneurons. To this end, we bred *Lhx6-Cre* mice with mice carrying floxed *Mef2c* alleles and with *Ai9* reporter mice to label Cre-expressing cells. In contrast to conditional *Ppargc1a* mutants, we found that the loss of Mef2c decreases the density and alters the laminar distribution of MGE-derived interneurons (Figures S8E and S8F), suggesting that Mef2c plays a role in the generation, migration, or survival of PV+ interneurons. Nevertheless, we found that, within the surviving MGE-derived cells, mice lacking one or two *Mef2c* alleles exhibit a dose-dependent loss of PGC-1⍺ in MGE-derived interneurons, which is accompanied by a parallel reduction in PV levels (Figures S8E and S8F).

The previous experiments suggested that Mef2c is required to express *Ppargc1a* and may subsequently form a complex with PGC-1⍺ to regulate the terminal differentiation of PV+ interneurons in response to neuronal activity in the postnatal cerebral cortex. Motif enrichment analysis at the loci bound by ERRγ also showed Mef2c as a potential interactor (Figure S7I). To decipher how PGC-1⍺, ERRγ , and Mef2c may regulate the expression of PV-enriched genes, we performed luciferase experiments in vitro with the long (⍺1) and short (⍺4) brain isoforms of PGC-1⍺^68^. We assessed whether different combinations of PGC-1⍺, ERRγ, and Mef2c regulate the expression of *Kcna1*, *Bcan*, and *Cplx1* by binding at the promoter of these genes on specific DNA sequences identified by the PGC-1⍺ CUT&RUN experiments (Figure S9). We found that PGC-1⍺1, ERRγ, and Mef2c together promote the expression of *Kcna1*. Both PGC-1⍺ isoforms (with the long isoform consistently having the most vigorous effect) and ERRγ can promote the expression of *Cplx1* and *Bcan*, but only when binding to *Bcan* promoter 2 (Figure 6I). Interestingly, Mef2c can negatively regulate *Bcan* expression when binding at *Bcan* promoter 1, suggesting that PGC-1⍺1, ERRγ, and Mef2c do not always work cooperatively (Figure 6I). This is consistent with previous work suggesting that ERRγ can directly interact with the two isoforms of PGC-1⍺, whereas only the long isoform of PGC-1⍺ has a Mef2c interaction site^69^. Similarly, Mef2c reduces the strong activation of *Cplx1* expression by ERRγ and PGC-1⍺1 or PGC-1⍺4, but it does not inhibit *Cplx1* expression alone or with PGC-1⍺. These results demonstrate that PGC-1⍺, through its interaction with ERRγ, directly controls the expression of genes required for the maturation of PV+ interneurons.

### PGC-1α is required for PV+ interneuron subtype specification

Several different subtypes of PV+ interneurons segregate across different layers of the cortex. Although they share many common features, each subtype of PV+ interneuron has specific molecular, electrophysiological, and morphological characteristics^70^. It has been proposed that neuronal activity may instruct the identity of cortical interneurons^4^, but this hypothesis has not been validated. Since *Ppargc1a* regulates the maturation of PV+ interneurons in an activity-dependent manner, we explored whether this gene may also contribute to the diversification of PV+ cells into different subtypes. To this end, we performed label transfer from a reference adult cortical interneuron dataset^70^ to identify layer-specific subtypes of PV+ interneurons in our dataset (Figure 7A). We found that the lack of PGC-1⍺ caused a shift in the proportion of PV+ interneuron subtypes, with an overrepresentation of L5/6 PV+ cells at the expense of L4 PV+ interneurons (Figure 7B). Although L4 PV+ interneurons can overcome the lack of PGC-1⍺ over time and acquire the general properties of PV+ cells, this observation suggests that some of these cells may do so at the expense of their specific L4 identity.

**Figure 7.**
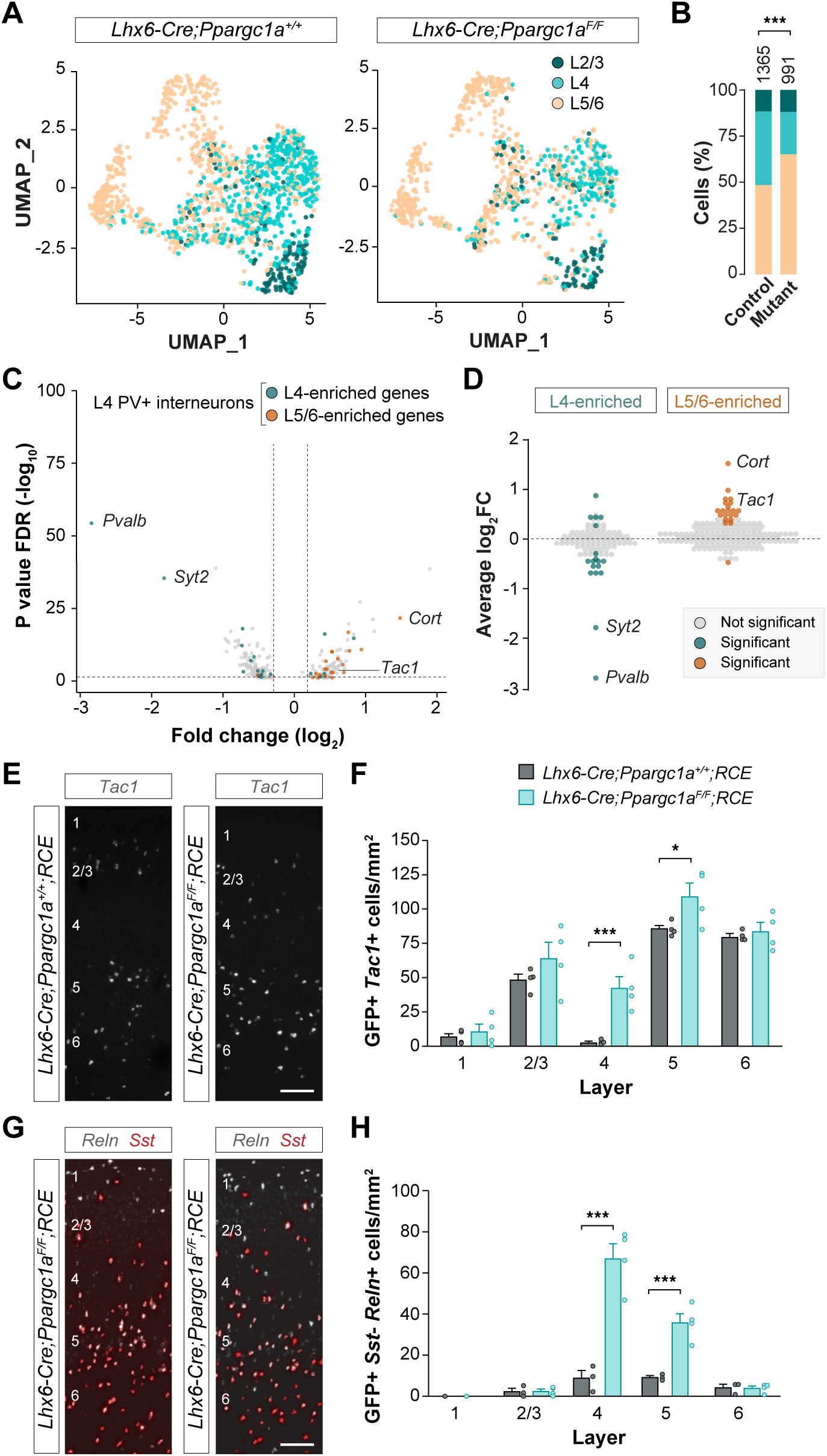
PGC-1⍺ regulates subtype specification in PV+ interneurons. (A) UMAP plots illustrating the assignment of L2/3, L4, and L5/6 subtypes of PV+ interneurons in control and conditional homozygous *Ppargc1a* mice at P21. (B) Proportion of PV+ interneurons identified as L2/3, L4, and L5/6 subtypes in control and conditional homozygous *Ppargc1a* mice at P21. Chi-square test, \*\*\**p* < 0.001. (C) Volcano plot illustrating DEGs between L4 PV+ interneurons in control and conditional homozygous *Ppargc1a* mice at P21. A threshold at log_2_FC > ±0.25 was set. Genes normally enriched in L4 and L5/6 PV+ interneurons are highlighted in cyan and orange, respectively. (D) Quantification of expression changes (average log_2_FC) in L4 PV+ interneurons between control and conditional homozygous *Ppargc1a* mice at P21 for genes normally enriched in L4 or L5/6 PV+ interneurons. A threshold at log_2_FC > ±0.25 was set. Significant changes in expression are highlighted in cyan and orange for genes normally enriched in L4 and L5/6 PV+ interneurons, respectively. (E) *Tac1* expression in coronal sections through S1 of control and conditional homozygous *Ppargc1a* mice at P21. (F) Quantification of the density of GFP+ *Tac1*+ cells in cortical layers of S1 in control and conditional homozygous *Ppargc1a* mice at P21. Two-way ANOVA followed by Tukey’s multiple correction tests, \**p* < 0.05, \*\*\**p* < 0.001. (G) *Reln* expression in PV+ interneurons (recognized by the absence of *Sst* expression) in coronal sections through S1 of control and conditional homozygous *Ppargc1a* mice at P21. (H) Quantification of the density of GFP+ *Sst*-*Reln*+ cells in cortical layers of S1 in control and conditional homozygous *Ppargc1a* mice at P21. Two-way ANOVA followed by Tukey’s multiple correction tests, \*\*\**p* < 0.001. Data are presented as mean ± s.e.m. Scale bars, 100 µm (E, G).

To investigate whether the remaining mutant L4 PV+ cells were abnormally specified, we performed a DGE analysis between L4 and L5/6 control PV+ interneurons at P21 and examined the expression of these genes in L4 PV+ interneurons lacking PGC-1⍺ (Figures 7C and 7D). We found that genes characteristic of L5/6 PV+ interneurons, such as *Tac1* and *Cort*, are abnormally upregulated in L4 PV+ interneurons in conditional *Ppargc1a* mutants (Figures 7C– 7F). Since *Tac1* and *Cort* are also expressed in L2/3 PV+ interneurons, we also examined the expression of *Reln*, enriched in L5 PV+ interneurons^70^. We found that *Reln* is also abnormally upregulated in L4 PV+ interneurons in conditional *Ppargc1a* mutants (Figures 7G and 7H), suggesting that PGC-1⍺ is required to specify L4 PV+ interneurons. Altogether, these observations suggest that PGC-1⍺ is not only required for the terminal differentiation of most PV+ interneurons but also seems to contribute to the diversification of PV+ interneuron subtypes in the early postnatal cortex.

## DISCUSSION

The mammalian cerebral cortex exhibits a characteristic protracted development. Cortical neurons are specified at embryonic stages but only acquire their adult features at relatively advanced stages of postnatal development. Here, we sought to identify the molecular mechanisms controlling the terminal differentiation of PV+ interneurons, a major group of GABAergic cells in the cerebral cortex. We found that expression of the transcriptional cofactor PGC-1⍺ is induced towards the end of the first week of postnatal development in cortical PV+ interneurons, where it functions as a molecular switch driving the transcriptional programs required for the maturation of these cells. PGC-1⍺ expression in PV+ interneurons is regulated by neuronal activity, thereby linking cortical dynamics to the timing of maturation of this critical population of inhibitory interneurons.

### Activity-dependent maturation of cortical PV+ interneurons

PV+ and SST+ interneurons constitute two of the main subclasses of interneurons in the mammalian cerebral cortex^1–3^. They both derive from progenitor cells in the MGE, populate the same layers of the neocortex, and are both reciprocally connected with pyramidal cells. However, the developmental trajectories of both subclasses of interneurons are asynchronous. SST+ interneurons are born earlier than PV+ interneurons and begin acquiring their mature morphological, electrophysiological, and transcriptomic features during the first week of postnatal development in mice^71,72^. This is substantially earlier than PV+ interneurons, which initiate their terminal differentiation well into the second week of postnatal development^28,73,74^. Our experiments revealed that the maturation of PV+ interneurons critically depends on neuronal activity. PV+ interneurons receive transient inhibitory connections from SST+ interneurons during the first week of postnatal development^75–77^, which effectively prevent their maturation during this period^24^. The number and strength of glutamatergic inputs received by PV+ interneurons increase progressively during the second week of postnatal development^28,74,75^, and this shift in balance between excitation and inhibition in PV+ interneurons seems to trigger the terminal differentiation of these cells. It remains to be determined whether a certain level of activity or a particular pattern of activity is required to drive the terminal differentiation of PV+ interneurons. Previous work has shown that the survival of cortical interneurons during the period of programmed cell death also depends on glutamatergic inputs^46–48^, and the pattern of activity seems critical in this context^78^.

We identified PGC-1⍺ as a master regulator of the terminal differentiation of PV+ interneurons, controlling the expression of a PV-specific transcriptional program. However, the precise molecular mechanisms leading to the onset of PGC-1⍺ expression in PV+ interneurons in response to activity remain unknown. Experiments in neuronal cultures indicated that PGC-1⍺ expression is modulated by Ca^2+^, AMK, and p38MAPK signaling, likely by recruiting the cAMP response element-binding protein (CREB) to the *Ppargc1a* locus^79–81^. The expression of MEF2 transcription factors is also promoted by glutamatergic signaling in neuronal cultures^82^. Interestingly, MEF2 transcription factors cooperate with CREB to regulate the expression of PGC-1⍺ in skeletal muscle^83^, and our experiments indicate that Mef2c is required in vivo for the expression of PGC-1⍺ in developing PV+ interneurons. However, Mef2c is already expressed by putative PV+ interneurons at embryonic stages^84^, long before *Ppargc1a* is expressed in these cells. This suggests that additional mechanisms contribute to the expression of PGC-1⍺ in PV+ interneurons. For instance, neuronal activity could induce activating post-translational modifications on Mef2c^85^ or induce epigenetic changes that contribute to the regulation of PGC-1⍺ expression, as shown in other tissues^86–88^.

### PGC-1α translates activity signals into specific transcriptional programs

Our experiments revealed that PGC-1⍺ is required to mediate the effect of neuronal activity in the maturation of PV+ interneurons. In other organs, PGC-1⍺ coordinates cellular responses to environmental cues^35,36^. For instance, PGC-1⍺ expression is induced in response to cold in skeletal muscle and brown fat adipose tissue, where it mediates adaptive thermogenesis^57,89^. In contrast, PGC-1⍺ expression is induced by fasting in the liver to promote gluconeogenesis^58,90,91^. In the brain, this transcriptional activator seems to have been coopted by cortical PV+ interneurons to link their terminal differentiation to the increase in neuronal activity that characterizes the second postnatal week of postnatal development.

PGC-1⍺ function in skeletal muscle, brown fat, and liver has been primarily linked to mitochondrial biogenesis and metabolism^57,92,93^. Analysis of mutant mice revealed that PGC-1α is dispensable for mitochondrial biogenesis per se, but it is required to express many mitochondrial genes in response to environmental cues^58,94^. We have found that PGC-1⍺ is also dispensable for mitochondrial biogenesis in PV+ and SST+ interneurons. However, we observed that PGC-1⍺ is critical for the dramatic surge in mitochondria that characterizes adult PV+ interneurons^54–56^ and for the expression of specific mitochondrial genes in these cells. This observation suggests that PGC-1⍺ drives the acquisition of the mature electrophysiological properties of PV+ interneurons while simultaneously promoting mitochondrial biogenesis in these cells, ensuring they can cope with the increased energy demand necessary to sustain their fast-spiking firing^10^.

Recent work suggests that mitochondrial metabolism serves as a pacemaker of neuronal maturation^60^. From this perspective, PGC-1⍺ could simply trigger the maturation of PV+ interneurons by promoting mitochondria biogenesis and oxidative phosphorylation in these cells. For instance, PGC-1⍺ may regulate the expression of enzymes that produce metabolites such as acetyl-CoA, critical for epigenetic regulation^95^. Consistently, conditional deletion of the cytochrome C subunits Cox10 or Cox6a2 from PV+ interneurons leads to a reduction in the levels of PV and perineuronal nets in these cells^96,97^. However, these defects do not match the severity of phenotypes observed in conditional *Ppargc1a* mutant mice. Moreover, our experiments reveal that PGC-1⍺ directly regulates the transcription of non-mitochondrial genes in PV+ interneurons, including voltage-dependent channels critical for their adult function. Consistent with this notion, PGC-1⍺ regulates the expression of cell type-specific programs in skeletal muscle, promoting the differentiation of slow-twitch muscle fibers^98^. Thus, it seems that PGC-1⍺ functions in the brain as a master regulator of PV+ interneurons, promoting mitochondrial biogenesis in parallel to cell type-specific transcriptional programs driving the terminal differentiation of these cells.

We found that PGC-1⍺ controls the expression of its brain-specific isoforms in cortical neurons. In addition, PGC-1⍺ interacts with ERRγ, whose expression is also directly controlled by PGC-1⍺, to regulate the expression of PV-enriched genes. The long isoform of PGC-1⍺ seems most critical for this function, although its short isoform is more highly expressed in S1 at P12. It is conceivable that long and short isoforms of PGC-1⍺ regulate different aspects of the maturation of PV+ interneurons because only the short isoform can enter the mitochondria and interact with mitochondrial DNA^93^. It has been previously suggested that ERR⍺, another member of the ERR family, is involved in the maturation of PV+ interneurons^65^. However, ERR⍺ is expressed at relatively low levels in the cerebral cortex and is not specific to PV+ interneurons, suggesting that ERRγ is the preferred coactivator of PGC-1⍺ in these cells. Moreover, ERRγ is only expressed in a small fraction of SST+ interneurons, which may explain the specificity of PGC-1⍺ in the differentiation of PV+ interneurons.

In addition to promoting the differentiation of PV+ interneurons through the expression of PV-specific genes, PGC-1⍺ and ERRγ seem to cooperate to repress the expression of SST-specific genes in these cells. This observation suggests that despite the early specification of PV+ and SST+ interneurons^66,99,100^, additional mechanisms contribute to consolidating the identity of these cells in the postnatal cortex. Consistent with this idea, we found that PGC-1⍺ modulates the specification of L4 PV+ interneurons. Compared to PV+ interneurons in other layers, L4 PV+ interneurons receive prominent inputs from the thalamus^101^. It is tempting to speculate that this additional source of activity may induce transcriptional programs that can compensate for the lack of PGC-1⍺ in these cells but may not be sufficient for the full acquisition of a specific L4 subtype identity. In sum, activity-dependent mechanisms are not only required for the terminal differentiation of PV+ interneurons but also contribute to their specification into distinct subtypes.

### Implications for neurodevelopmental disorders

The maturation of fast somatic inhibition by PV+ interneurons is critical for the emergence of sparse activity in cortical networks^24,102,103^, the regulation of critical periods of plasticity^25–27^, and the generation of gamma oscillations essential for information processing^14,15,104^. Unsurprisingly, abnormal development and dysfunction of cortical PV+ interneurons have been repeatedly linked with neurodevelopmental disorders^105,106^. For instance, postmortem studies have shown that PV expression in the cerebral cortex is consistently downregulated in autism spectrum disorder (ASD) and schizophrenia^107–111^, and mice carrying loss of function variants of developmental genes associated with these conditions exhibit reduced levels of PV in the cerebral cortex^112–116^. Although genetic variation in the *Ppargc1a* locus has not been directly linked to neurodevelopmental disorders, loss of function mutations in *MEF2C* are linked to intellectual disability, ASD, and schizophrenia^117^. Our study suggests that PV+ interneurons are particularly susceptible to network dynamics to conclude their differentiation, which is crucial for cortical function. For example, several genes whose variation have been linked to neurodevelopmental disorders regulate the development of excitatory synapses on PV+ interneurons^42,118,119^, and these early wiring defects may impair PGC-1⍺ expression and the maturation of these cells. In addition, PV+ interneurons are more susceptible to redox dysregulation and oxidative stress than other neurons due to their high metabolic demands^120^. Thus, defects in the maturation of PV+ interneurons may render these cells unable to cope with subsequent environmental insults, leading to cortical dysfunction and disease.

## ACKNOWLEDGEMENTS

We thank E. Serafeimidou-Pouliou and C. Zimmer for general laboratory support, I. Andrew for managing mouse colonies, and the Flow Cytometry and Genomics Research Platforms at King’s College London for the scRNA-seq experiments. We are also grateful to B. Rico and T. Garcés for the E2-Cre plasmid, P. Vanderhaeghen and R. Iwata for the mito-EmGFP plasmid, and members of the Marín and Rico laboratories for stimulating discussions and ideas. This work was supported by grants from the Wellcome Trust (215556/Z/19/Z) to O.M.

## AUTHOR CONTRIBUTIONS

Conceptualization, M.M. & O.M.; Methodology, M.M., L.A., R.A., C.B., Y.D., S.Q., A.K., L.M., & F.H.; Investigation, M.M., R.A., C.B., Y.D., & A.K.; Data curation, M.M., L.A., R.A., C.B., & P.L.; Formal Analysis, M.M., L.A., R.A., C.B., & F.H.; Writing – Original Draft, M.M. & O.M.; Writing – Review & Editing, M.M., P.L., N.F., & O.M.; Funding Acquisition, O.M.; Resources, P.L., N.F. & O.M.; Supervision, P.L., N.F. & O.M.

---

**Figure S1.**
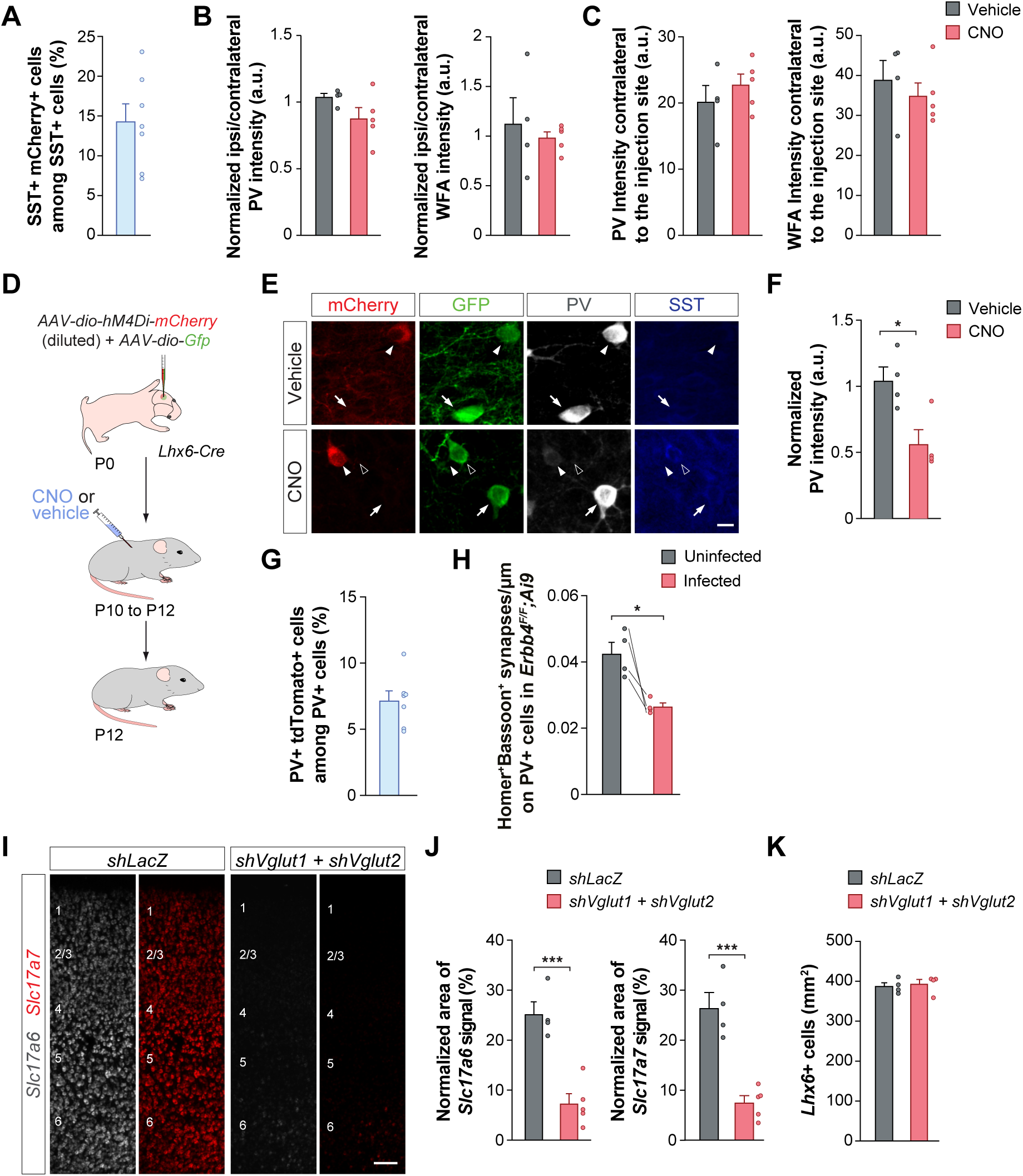
The maturation of cortical PV+ interneurons depends on neuronal activity. (A) Infection rate of S1 L2/3 SST+ cells by *AAV8-hSyn-dio-hM3Dq-mCherry*. (B) Quantification of detectable PV and WFA levels in S1 L2/3 PV+ interneurons of the ipsilateral side of injection, normalized to detectable PV and WFA levels of the contralateral side of injection, in control (vehicle) and experimental (CNO) conditions at P12. Unpaired Student’s *t*-test, not significant. (C) Quantification of detectable PV and WFA levels in S1 L2/3 PV+ interneurons of the contralateral side of injection in control (vehicle) and experimental (CNO) conditions at P12. Unpaired Student’s *t*-test, not significant. (D) Schematic representation of the experimental design. (E) PV expression in S1 L5 prospective PV+ cells (recognized by the absence of SST expression) expressing mCherry or GFP in control (vehicle) and experimental (CNO) conditions at P12. Arrows point at GFP+ PV+ cells, arrowheads point at mCherry+ PV+ cells, and open arrowheads point at GFP+ SST+ cells. (F) Quantification of PV levels in S1 L5 mCherry+ prospective PV+ cells normalized by PV levels in neighboring mCherry-GFP+ prospective PV+ cells at P12. Unpaired Student’s *t*-test, \**p* < 0.05. (G) Infection rate of S1 L5/6 PV+ cells by the *AAV8-E2-Cre* virus. (H) Quantification of the density of Homer+ Bassoon+ synapses contacting the soma of tdTomato+ PV+ cells compared to neighboring uninfected PV+ cells in *ErbB4^F/F^;Ai9* mice at P12. (I) *Slc7a6* and *Slc7a7* expression in coronal sections through S1 from control (*shLacZ*) and experimental (*shVglut1 + shVglut2*) mice at P12. (J) Quantification of the area occupied by *Slc7a6* or *Slc7a7* normalized to the total area size of the image in S1 from control (*shLacZ*) and experimental (*shVglut1 + shVglut2*) mice at P12. Unpaired Student’s *t*-test, \*\*\**p* < 0.001. (K) Quantification of the density of *Lhx6*+ cells in S1 from control (*shLacZ*) and experimental (*shVglut1 + shVglut2*) mice at P12. Unpaired Student’s *t*-test, not significant. Data are presented as mean ± s.e.m. Scale bars, 10 µm (E), 100 µm (G).

**Figure S2.**
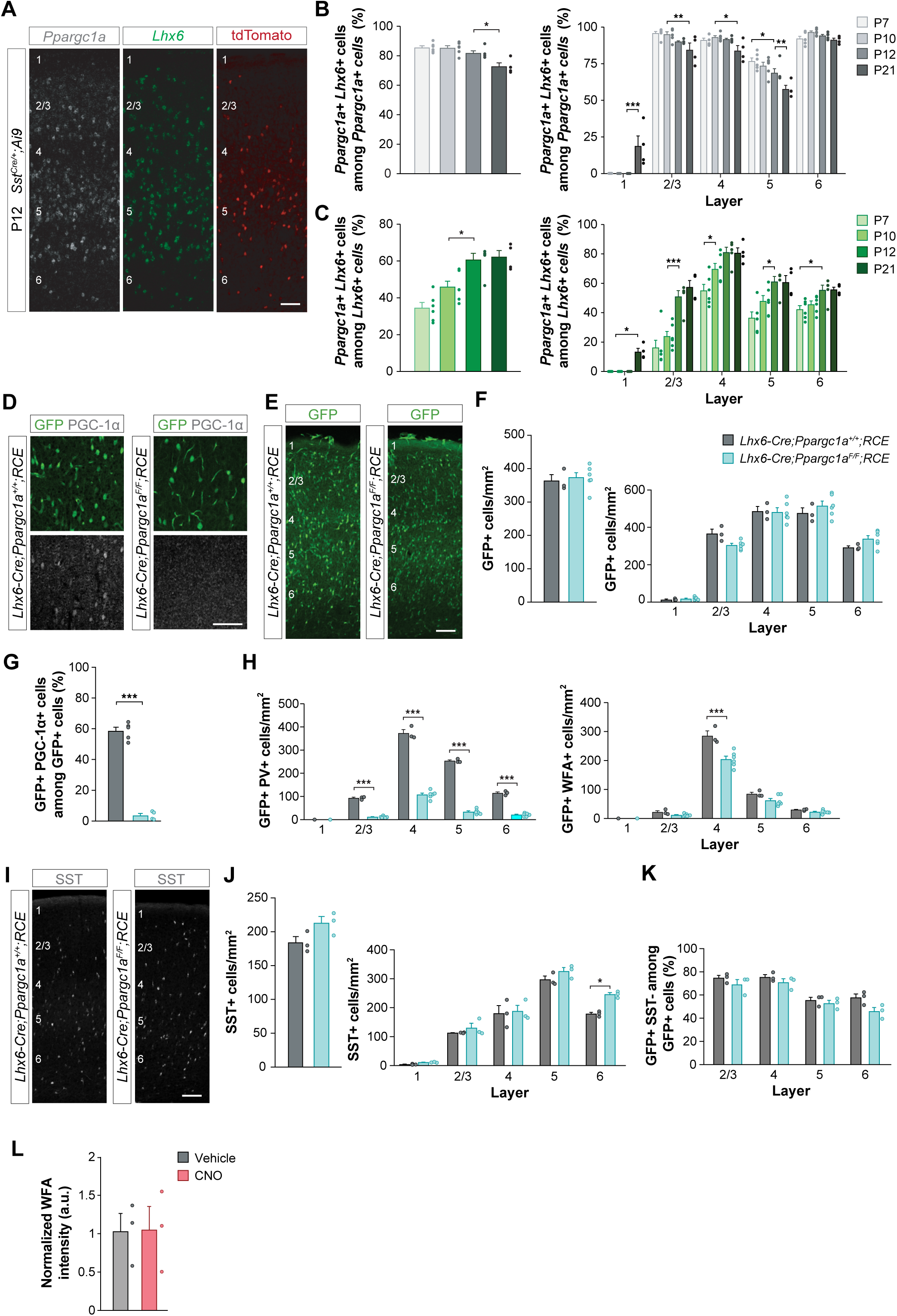
The onset of *Ppargc1a* expression precedes the onset of maturation of cortical PV+ interneurons, which depends on the activity-dependent expression of PGC-1⍺. (A) *Ppargc1a*, *Lhx6,* and tdTomato expression in coronal sections through S1 of *Sst^Cre/+^;Ai9* mice at P12. (B) Quantification of the proportion of *Ppargc1a*+ cells being *Lhx6*+ in all cortical layers at P7, P10, P12 and P21. One-way and two-way ANOVA followed by Tukey’s multiple correction tests, \**p* < 0.05, **p < 0.01, \*\*\**p* < 0.001. (C) Quantification of the proportion of *Lhx6*+ cells being *Ppargc1a*+ in all cortical layers at P7, P10, P12 and P21. One-way and two-way ANOVA followed by Tukey’s multiple correction tests, \**p* < 0.05, **p < 0.01, \*\*\**p* < 0.001. (D) PGC-1⍺ expression in GFP+ cells in S1 of control and conditional homozygous *Ppargc1a* mice at P21. (E) GFP expression in coronal sections through S1 of control and conditional homozygous *Ppargc1a* mice at P12. (F) Quantification of the density of GFP+ cells in S1 of control and conditional homozygous *Ppargc1a* mice at P12. One-way and two-way ANOVA followed by Tukey’s multiple correction tests, not significant. (G) Quantification of the proportion of GFP+ cells being PGC-1⍺+ in S1 of control and conditional homozygous *Ppargc1a* mice at P21. Unpaired Student’s *t*-test, \*\*\**p* < 0.001. (H) Quantification of the density of GFP+ PV+ cells and GFP+ WFA+ cells in cortical layers of S1 in control and conditional homozygous *Ppargc1a* mice at P12. Two-way ANOVA followed by Tukey’s multiple correction tests, \*\*\**p* < 0.001. (I) SST expression in coronal sections through S1 of control and conditional homozygous *Ppargc1a* mice at P12. (J) Quantification of the density of GFP+ SST+ cells in cortical layers of S1 in control and conditional homozygous *Ppargc1a* mice at P12. Two-way ANOVA followed by Sidak’s multiple correction tests, \**p* < 0.05. (K) Quantification of the proportion of GFP+ SST- (i.e., prospective PV+ interneurons) in cortical layers of S1 in control and conditional homozygous *Ppargc1a* mice at P12. Two-way ANOVA followed by Sidak’s multiple correction tests, not significant. (L) Quantification of WFA levels in S1 L2/3 mCherry+ prospective PV+ cells normalized by WFA levels in neighboring mCherry-GFP+ prospective PV+ cells at P12. Unpaired Student’s *t*-test, not significant. Data are presented as mean ± s.e.m. Scale bars, 100 µm (A, D, E, and I)

**Figure S3.**
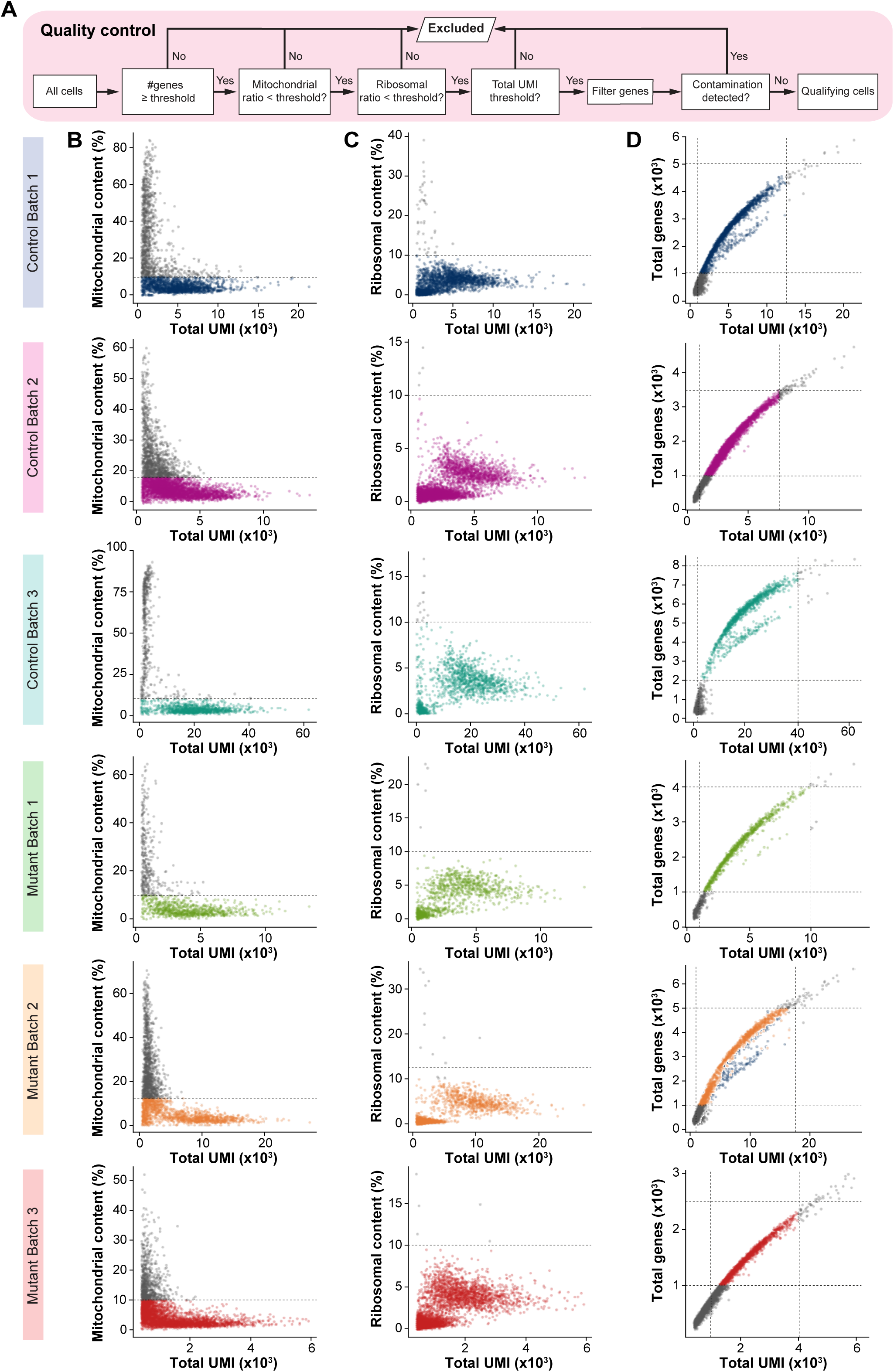
Quality control in single-cell RNA sequencing experiments. (A) Schematic representation of the quality control pipeline. (B, C) Percentage of mitochondrial and ribosomal content in each cell in each control and mutant batches of cells. (D) Total number of genes detected in each cell in each control and mutant batches.

**Figure S4.**
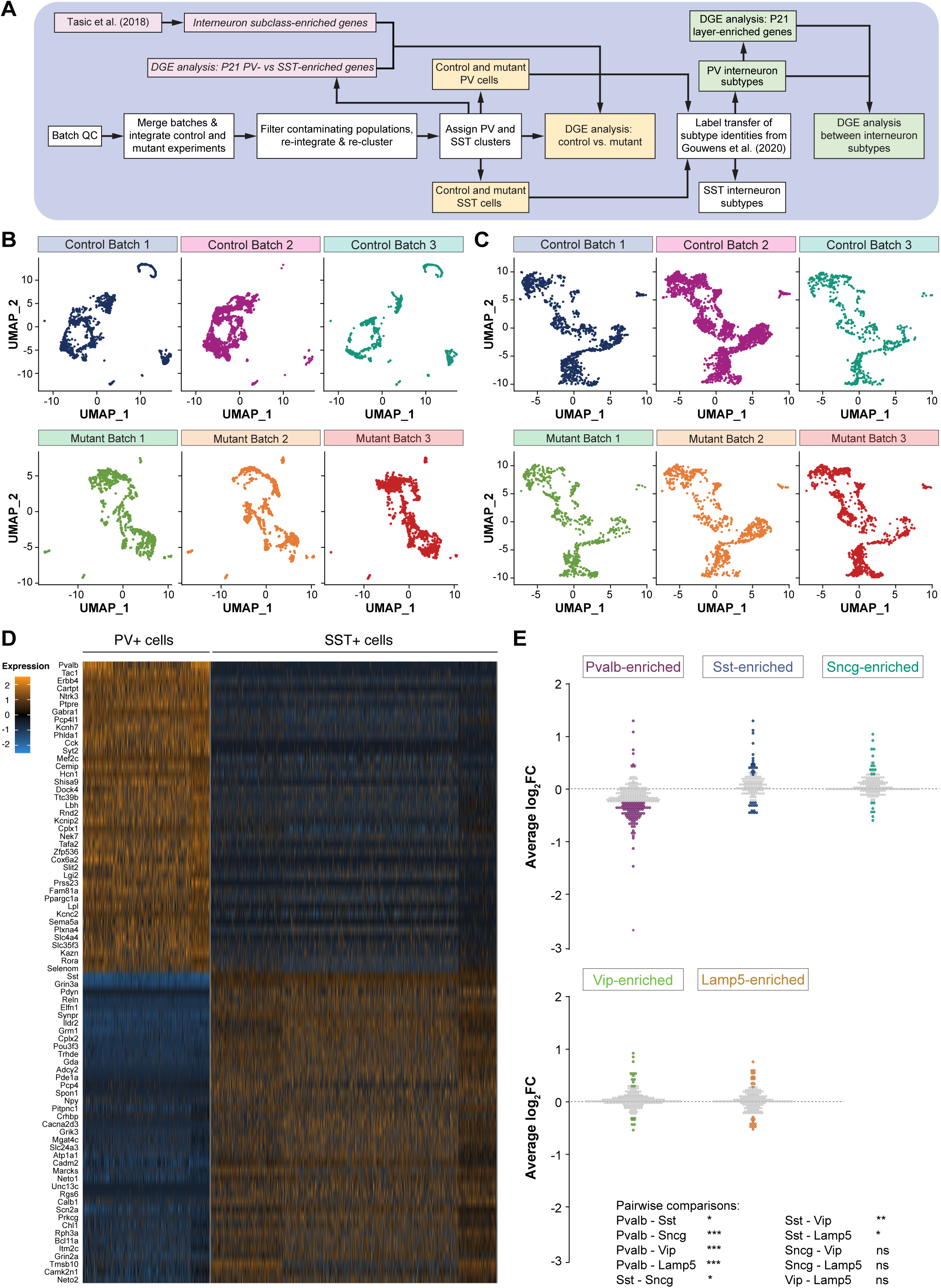
Gene expression alterations in PV+ and SST+ interneurons lacking PGC-1⍺. (A) Schematic representation of the analysis pipeline following single-cell RNA sequencing of control and conditional homozygous *Ppargc1a* mice at P21. (B) UMAP plots depicting the cells sequenced in the different control and mutant batches before integration of all batches. (C) UMAP plots depicting the cells sequenced in the different control and mutant batches after integration of all batches. (D) Heatmap representing differential gene expression between control PV+ and SST+ interneurons at P21. Only the top 40 genes are represented. (E) Quantification of expression changes (average log_2_FC) in PV+ interneurons between control and conditional homozygous *Ppargc1a* mice at P21 for genes normally enriched in the five main subclasses of cortical interneurons. A threshold at log_2_FC > ±0.25 was set. Significant changes in expression are highlighted in magenta, dark blue, dark green, green, and orange for genes normally enriched in Pvalb, Sst, Sncg, Vip, and Lamp5 subclasses of interneurons, respectively. Fisher’s exact test, \**p* < 0.05, \*\*\**p* < 0.001.

**Figure S5.**
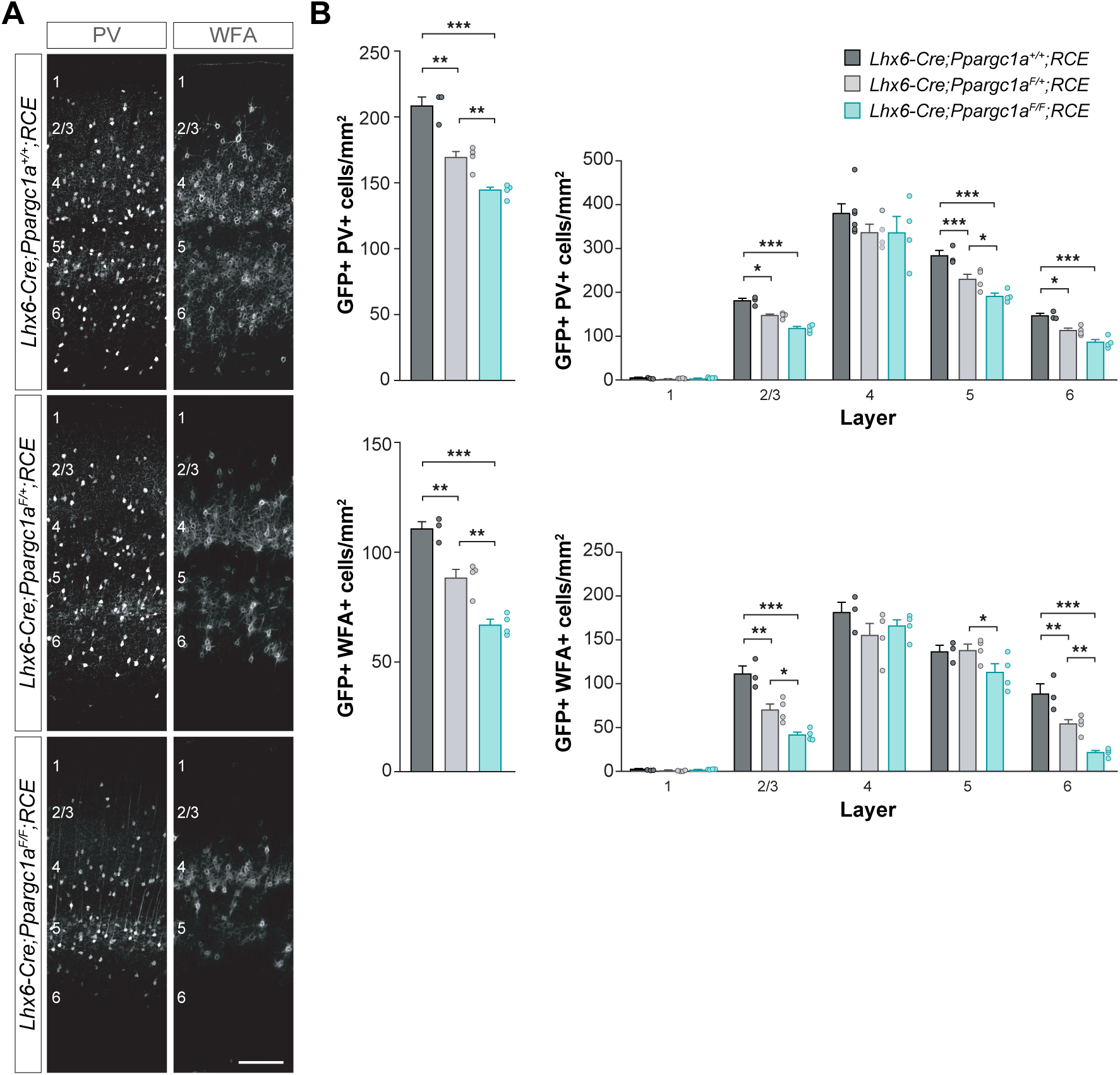
Loss of PGC-1⍺ halts the maturation of many cortical PV+ interneurons. (A) PV and WFA expression in coronal sections through S1 of control and conditional homozygous *Ppargc1a* mice at P60. (B) Quantification of the density of GFP+ PV+ cells and GFP+ WFA+ cells in S1 of control and conditional homozygous *Ppargc1a* mice at P60. One-way and two-way ANOVA followed by Tukey’s multiple correction tests, \**p* < 0.05, \*\**p* < 0.01, \*\*\**p* < 0.001. Data are presented as mean ± s.e.m. Scale bar, 100 µm (A)

**Figure S6.**
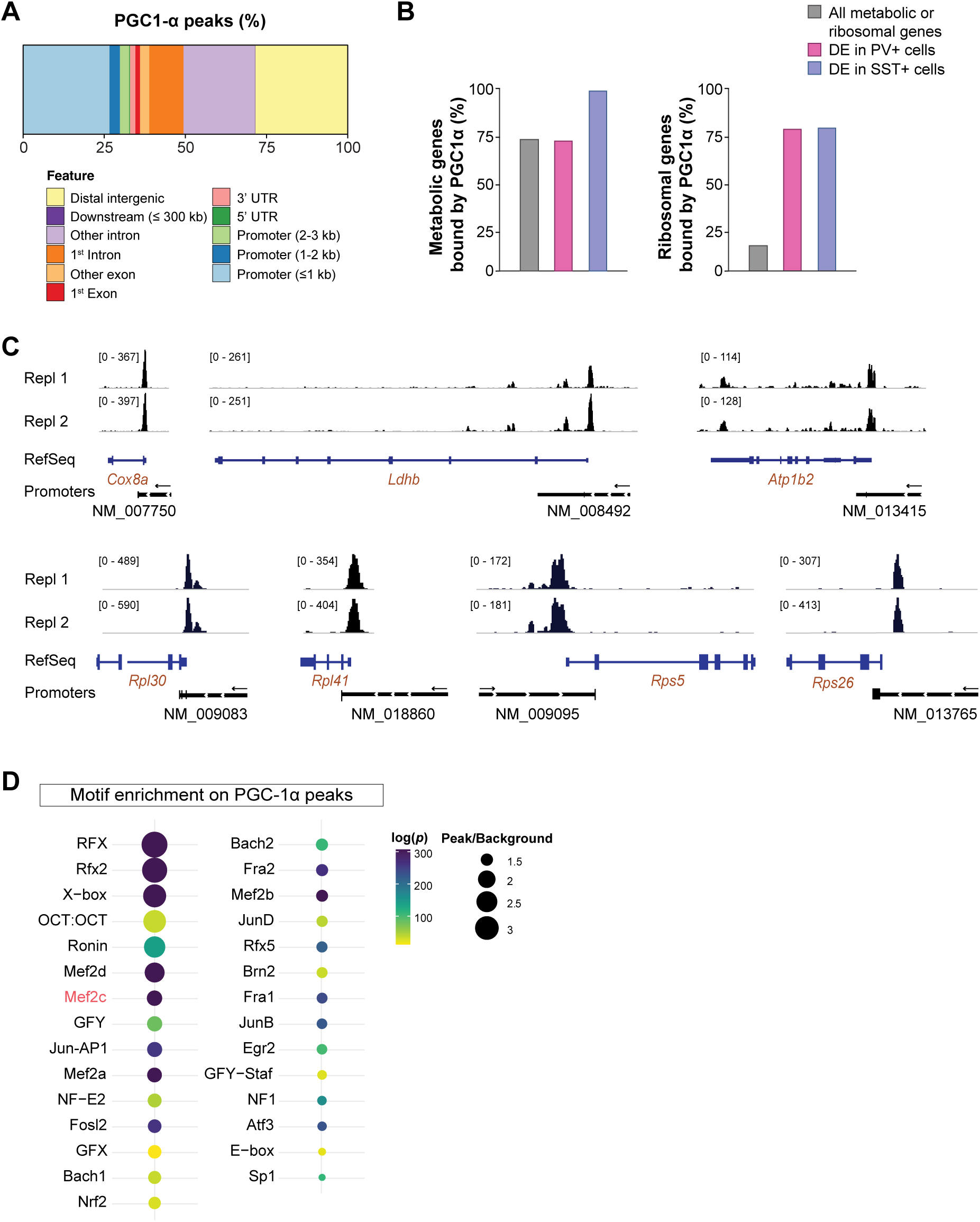
PGC-1⍺ regulates gene expression through direct transcriptomic regulation. (A) Proportion of PGC-1⍺ peaks observed on the different gene features (distal intergenic, introns, exons, UTR, promoters). (B) Plots depicting the proportion of all metabolic or all ribosomal genes directly bound by PGC-1⍺ (grey) and the proportion of metabolic or ribosomal DEGs between control and conditional homozygous *Ppargc1a* PV+ cells (magenta) and SST+ cells (blue) directly bound by PGC-1⍺. (C) Genome browser view of the genomic locus of metabolic (*Cox8a, Ldhb,* and *Atp1b2*) and ribosomal (*Rpl30, Rpl41, Rps5,* and *Rps26*) genes. The black peaks represent the read coverage following PGC-1⍺ CUT&RUN in two replicates at P12. Gene promoters, and the size scale of the peaks, are annotated. The arrows above the promoters indicate the direction of transcription. (D) Motif enrichment analysis at the site of PGC-1⍺ peaks, revealing potential binding partners of PGC-1⍺ to regulate gene expression. Peak/background represents the incidence of the motif among target peaks over the incidence among background sequences.

**Figure S7.**
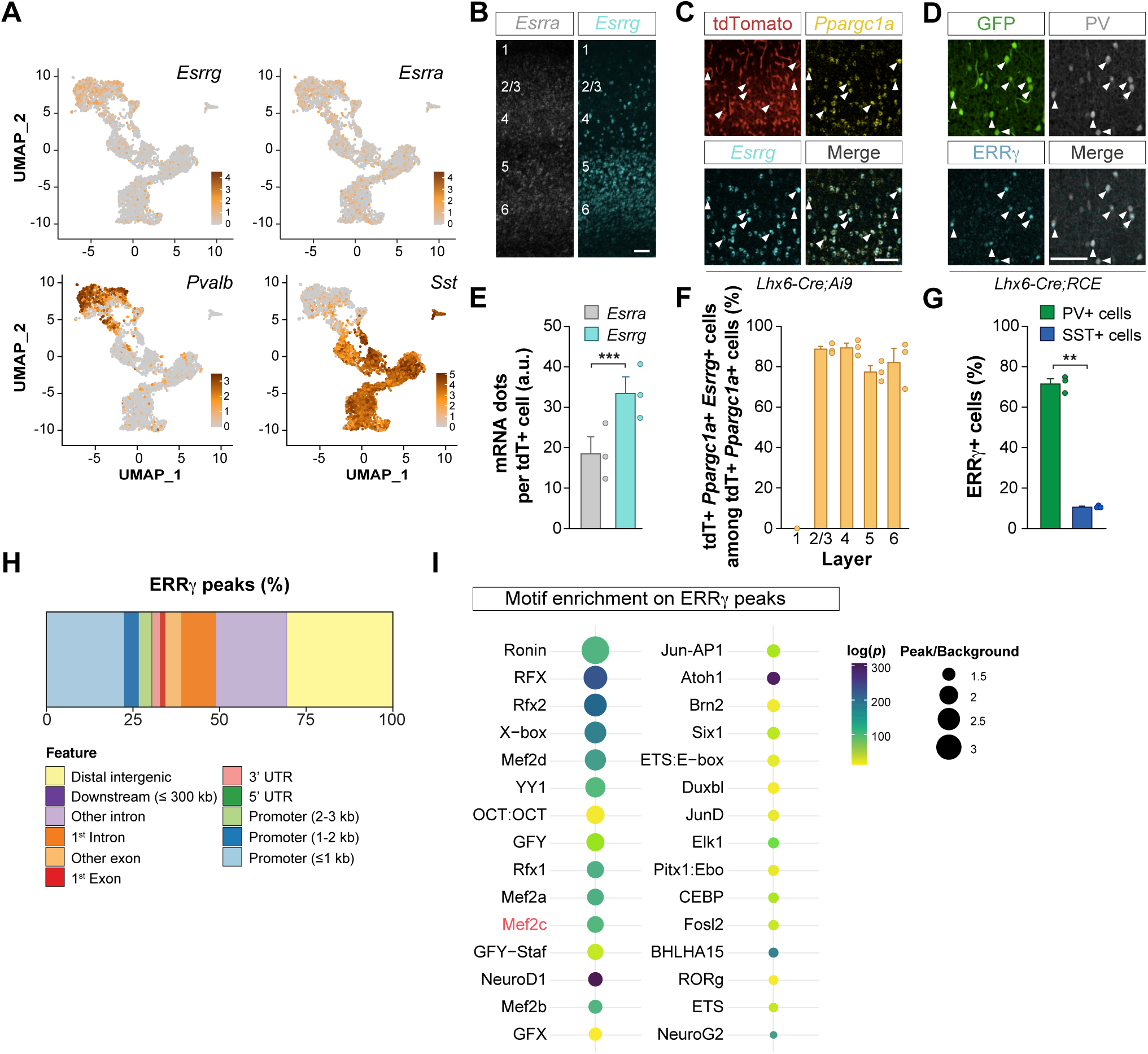
Expression of *Esrrg* and ERRγ in MGE-derived interneurons and analysis of ERRγ genomic binding sites. (A) UMAP plots illustrating the expression of *Esrrg*, *Esrra*, *Pvalb,* and *Sst* in MGE-derived cells at P21. (B) *Esrra* and *Esrrg* expression in coronal sections through S1 of *Lhx6-Cre;Ai9* mice at P12. (C) *Esrrg* and *Ppargc1a* expression in tdTomato+ cells in S1 of *Lhx6-Cre;Ai9* mice at P12. (D) ERRγ expression in GFP+ PV+ interneurons and prospective SST+ (GFP+ PV-) interneurons in S1 of *Lhx6-Cre;RCE* mice at P21. (E) Quantification of the number of *Esrra* and *Esrrg* RNAscope dots present in tdTomato+ cells in S1 L2-4 of *Lhx6-Cre;Ai9* mice at P12. Unpaired Student’s *t*-test, \*\*\**p* < 0.001. (F) Quantification of the proportion of tdTomato+ *Ppargc1a*+ cells expressing *Esrrg* at P12 through cortical layers of S1 of *Lhx6-Cre;Ai9* mice. (G) Quantification of the proportion of PV+ and prospective SST+ cells expressing ERRγ in S1 of *Lhx6-Cre;RCE* animals at P21. Unpaired Student’s *t*-test, \*\**p* < 0.01. (H) Proportion of ERRγ peaks observed on the different gene features (distal intergenic, introns, exons, UTR, promoters). (I) Motif enrichment analysis at the site of ERRγ peaks, revealing potential binding partners of ERRγ to regulate gene expression. Peak/background represents the incidence of the motif among target peaks over the incidence among background sequences. Data are presented as mean ± s.e.m. Scale bars, 10 µm (C, D), 100 µm (B).

**Figure S8.**
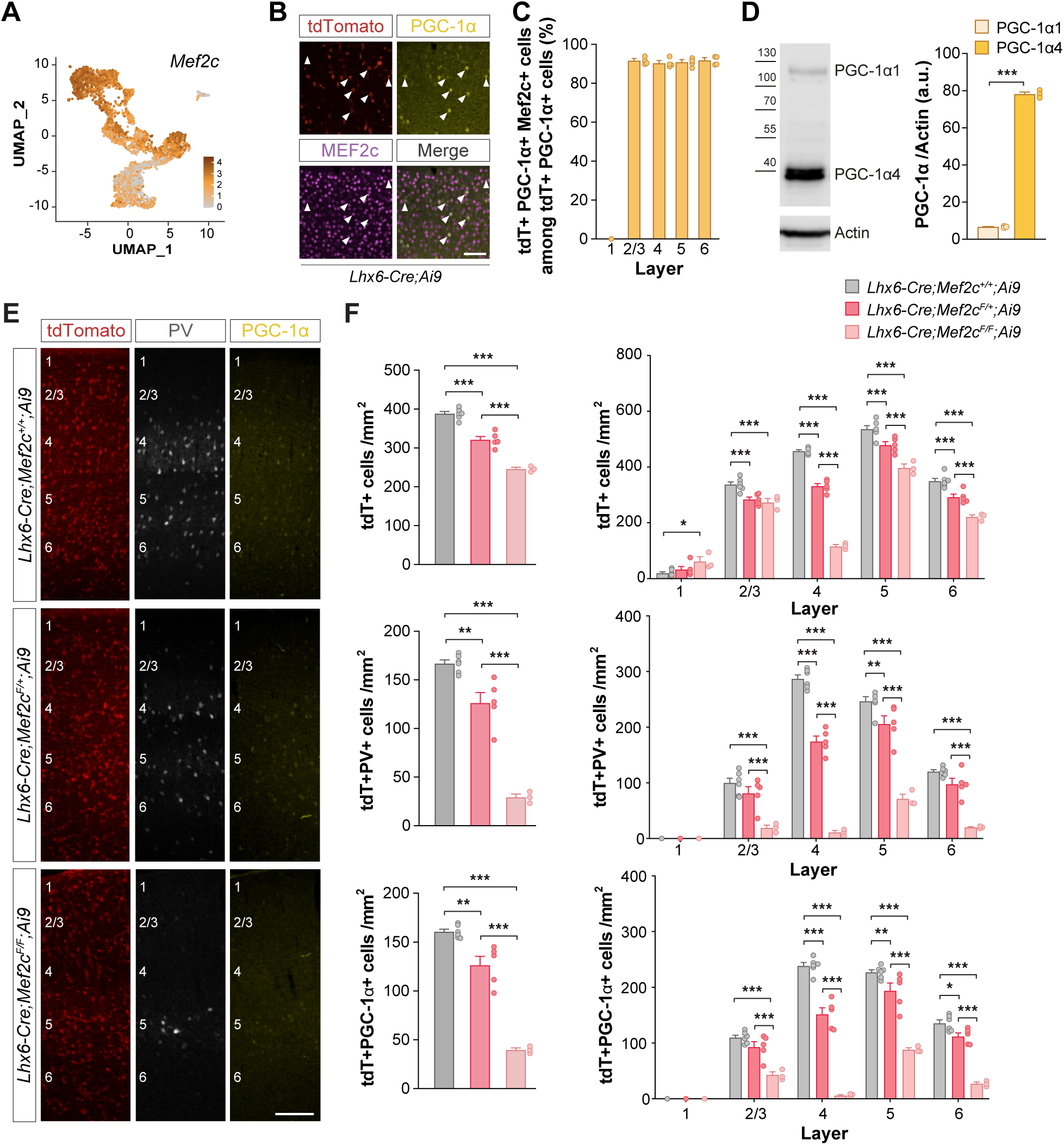
Mef2c is expressed in MGE-derived interneurons and is required for the expression of PGC-1⍺. (A) UMAP plot illustrating the expression of *Mef2c* in MGE-derived cells at P21. (B) Mef2c and PGC-1⍺ expression in tdTomato+ cells in S1 of *Lhx6-Cre;Ai9* animals at P12. (C) Quantification of the proportion of tdTomato+ PGC-1⍺ + cells expressing Mef2c at P12 through cortical layers of S1 in *Lhx6-Cre;Ai9* animals. (D) Western Blot image and quantification of the levels of expression of the short (PGC-1⍺4) and long (PGC-1⍺1) PGC-1⍺ isoforms in S1 at P12. (E) PV, PGC-1⍺, and tdTomato expression in coronal sections through S1 of control, conditional heterozygous, and conditional homozygous *Mef2c* mutant mice at P12. (F) Quantification of the density of tdTomato+ cells, tdTomato+ PV+ cells, and tdTomato+ PGC-1⍺+ cells in S1 of control, conditional heterozygous, and conditional homozygous *Mef2c* mutant mice at P12 across all cortical layers. One-way and two-way ANOVA followed by Tukey’s multiple correction tests, \**p* < 0.05, \*\**p* < 0.01, \*\*\**p* < 0.001. Data are presented as mean ± s.e.m. Scale bars, 10 µm (B), 100 µm (E).

**Figure S9.**
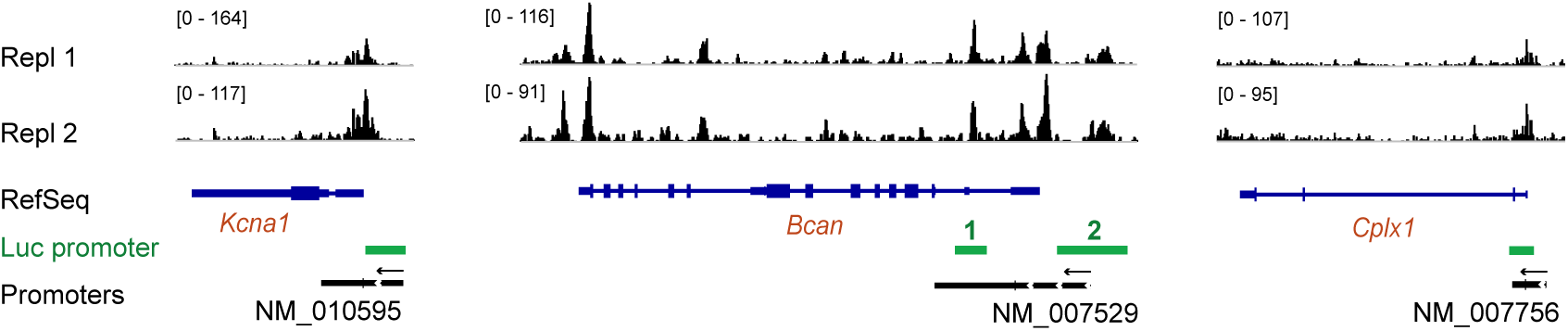
Characterization of luciferase assay experiments. Genome browser view of the genomic locus of *Kcna1*, *Bcan*, and *Cplx1*. The black peaks represent the read coverage following PGC-1⍺ CUT&RUN in two replicates at P12. Gene promoters (in black) and promoters chosen to conduct the luciferase assays (in green), and the size scale of the peaks, are annotated. The arrows above the promoters indicate the direction of transcription.

## METHODS

### Mice

All experiments were performed following King’s College London Biological Service Unit’s guidelines and under license from the UK Home Office following the Animals (Scientific Procedures) Act 1986. Similar numbers of male and female mice were used in all experiments. Animals were maintained under standard laboratory conditions on a 12:12 light/dark cycle with water and food ad libitum. We used *Lhx6-Cre* mice [B6;CBA-Tg(Lhx6-icre)1Kess/J]^50^ to modulate the activity of PV+ interneurons at early postnatal stages. *ErbB4*-floxed mice [Erbb4^tm1Fej^]^123^ crossed to *Ai9* mice [Gt(ROSA)26Sor^tm9(CAG–tdTomato)Hze^]^122^ (JAX007909), and *Nex^Cre/+^* mice [Neurod6^tm1(cre)Kan^]^125^ were used to modulate the excitatory activity received by PV+ interneurons. To investigate the pattern of *Ppargc1a* and PGC-1⍺ expression, we crossed *Sst^Cre/+^* mice [Sst^tm2.1(cre)Zjh^]^127^ with *Ai9* mice, and *Lhx6-Cre* mice to *RCE* mice [Gt(ROSA)26Sor^tm1.1(CAG–EGFP)Fsh^]^126^ (JAX032037). To characterize the role of PGC-1⍺ on the maturation of PV+ interneurons, mice carrying loxP-flanked *Ppargc1a* alleles [Ppargc1a^tm2.1Brsp^]^58^ (JAX009666) were crossed to *Lhx6-Cre* or *Lhx6-Cre;RCE* mice. *Lhx6-Cre;Ai9* mice were used to assess the pattern of expression of MEF2c, *Esrra,* and *Esrrg*. *Mef2c*-floxed mice [Mef2c^tm1Jjs^]^124^ (JAX025556) were crossed to *Lhx6-Cre;Ai9* mice to study the role of Mef2c in the development of PV+ interneurons. WT C57BL6 mice (Charles River) were used for CUT&RUN experiments.

### Viral production

Serotype 8 adeno-associated viruses (AAV8) were produced in HEK293FT cells grown on 5 (Polysciences Europe, 23966-100) or 10 15-cm diameter plates (Sigma-Aldrich #408727), depending on whether we used linear or branched polyethylenimine (PEI) transfection, respectively, until cells reached 60% confluency. Cells were grown on DMEM (Gibco #21969-035) supplemented with 10% fetal bovine serum (FBS) (Gibco #10500-064), 1% penicillin/streptomycin (Gibco #15140-122), and 10 mM HEPES. AAVs were produced using PEI transfection of HEK293FT cells with a virus-specific transfer plasmid (70µg/10 plates) and a pDP8.ape helper plasmid (300 µg/10 plates; PlasmidFactory #PF478). The helper plasmid provided the AAV Rep and Cap functions and the Ad5 genes (VA RNAs, E2A, and E4). The DNA and PEI were mixed in a 1:4 ratio in uncomplemented DMEM and left at room temperature for 25 min to form the DNA-PEI complex. The transfection solution was added to each plate and incubated for 72h at 37°C in 5% CO_2_. The transfected cells were then scraped off the plates and pelleted. The cell pellet was lysed in buffer containing 50 mM Tris-Cl, 150 mM NaCl, 2 mM MgCl_2_, and 0.5% Sodium deoxycholate and incubated with 100 U/mL Benzonase nuclease (Sigma #E1014 25KU) for 1 h to dissociate particles from membranes. The particles were cleared by centrifugation, and the clear supernatant was filtered through 0.8 µm (Merck Millipore #SLAA033SS) and 0.45µm (Merck Millipore #SLHA 033SS) filters. The viral suspension was loaded on a discontinuous iodixanol gradient using four layers of different iodixanol concentrations^128^ of 15%, 25%, 40%, and 58% in Quick-seal polyallomer tubes (Beckman Coulter #342414) and spun in a Vti-50 rotor at 50.000 rpm for 75 min at 12°C in an Optima L-100 XP Beckman Coulter ultracentrifuge to remove any remaining contaminants. After completion of the centrifugation, 5mL was withdrawn from the 40-58% interface using a G20 needle. The recovered virus fraction was purified by first passing through a 100 kDa molecular weight cutoff (MWCO) centrifugal filter (Sartorius, VIVASPIN VS2041) and then through an Amicon Ultra 2mL Centrifugal filter (Millipore #UFC210024). Storage buffer (350 mM NaCl and 5% Sorbitol in PBS) was added to the purified virus, and 5 µl aliquots were stored at -80°C.

### Viral injections

Pups were separated from the dam and anesthetized with 1.5% isoflurane via a nose cone and placed in a stereotaxic frame containing a heating pad. Glass capillaries with a tip diameter of 30-45 µm were attached to a Nanoinjector (Drummond). Pups were injected with specific viral mixes, depending on the experiment, at Z-depth coordinates -0.2, -0.4, -0.6 to target cells of all cortical layers and delivered in two locations of S1 of the left hemisphere at an injection rate of 5nL/s (unless specified otherwise below). Pups were allowed to recover in a heating chamber at 37°C and then returned to the dam.

#### DREADD experiments

*Lhx6-Cre* or *Lhx6-Cre;Ppargc1a^F/F^*pups were injected at postnatal day (P)0 or P1 with 600nL of the following viral mix: *AAV8-hSyn-dio-hM3D(Gq)-mCherry* (Addgene #44361-AAV8) at a 1:40 dilution for sparse cell targeting, mixed with *AAV8-hSyn-dio-GFP* (Addgene #50457-AAV8, 1:2 dilution with sterile PBS and 0.5% Fast Green (Sigma)). *Lhx6-Cre* pups were also injected at P0 or P1 in a separate experiment, with 600 nL of a viral mix composed of *AAV8-hSyn-dio-hM4D(Gi)-mCherry* (Addgene #44362-AAV8) at a 1:40 dilution for sparse cell targeting and *AAV8-hSyn-dio-GFP* (Addgene #50457-AAV8, 1:2 dilution with sterile PBS and 0.5% Fast Green, Sigma). The dilution factor of the DREADDs viruses (here 1:40) was calculated to reach a titer of 8.10^11^ for sparse labeling.

#### Excitatory innervation experiments

To assess the role of excitatory innervation on the maturation of PV+ interneurons, *ErbB4^F/F^;Ai9* mice and *ErbB4^+/+^;Ai9* mice were injected at P1 with 600 nL of custom-made *AAV8-E2-Cre*, diluted at 1:15 with sterile PBS and 0.5% Fast Green (Sigma). The Rico lab gifted the E2-Cre construct, which was subsequently cloned into an AAV8 backbone. The dilution factor of the virus (here 1:15) was calculated to reach a titer of 8.10^11^ for sparse labeling. We observed leakage of Cre recombination in excitatory cells following injection at P1, which was ignored since ErbB4 is only expressed in putative PV interneurons at postnatal ages.

#### Glutamate release experiments

To assess the role of glutamate release by excitatory neurons on the maturation of PV+ interneurons, custom-made *AAV8-hSyn-dio-shLacZ* or a mix (1-to-1 ratio) of *AAV8-hSyn-dio-shVglut1* and *AAV8-hSyn-dio-shVglut2* colored with 0.5% Fast Green (Sigma) were injected at P6 in *Nex^Cre/+^* mice. We used previously designed shRNA sequences to target Vglut1 and Vglut2^129,130^. A volume of 600 nL was injected at a speed of 50 nL/min. The skin of the animals was first cut with a blade, the capillary containing the viral mix was pulled down through the soft skull at the coordinates specified above, and the skin was closed after surgery using surgical glue (3M Vetbond #1469SB).

#### Single-cell RNA-sequencing experiments

*Lhx6-Cre;Ppargc1a^+/+^*and *Lhx6-Cre;Ppargc1a^F/F^* animals were injected in both hemispheres at P1 with 900nL of *AAV8-hSyn-dio-tdTomato* colored with 0.5% Fast Green (Sigma) (300nL injected at three different locations of S1 in each hemisphere).

#### Mito-EmGFP experiments

*Lhx6-Cre;Ppargc1a^+/+^* and *Lhx6-Cre;Ppargc1a^F/F^* animals were injected at P1 with 600nL of custom-made *AAV8-hSyn-dio-mito-EmGFP* (diluted 1:2 with sterile PBS and 0.5% Fast Green, Sigma). The Vanderhaeghen lab gifted the mito-EmGFP construct, which was subsequently cloned into an AAV8 backbone.

### CNO administration

Clozapine-N-Oxide (CNO, Tocris #4936) was dissolved in 5% dimethyl sulfoxide (Sigma) and then diluted with 0.9% saline to 0.1mg/mL. Pups were injected intraperitoneally with vehicle (0.05% DMSO) or CNO (10 µL/g) twice daily for two days and a half (from P10 to P12). Vehicle and CNO treatments were randomly assigned among littermates.

### Histology

#### Immunohistochemistry

Mice were anesthetized with an overdose of sodium pentobarbital and transcardially perfused with saline, followed by 4% paraformaldehyde (PFA). Brains were postfixed 2h at 4°C, cryoprotected in 15% sucrose followed by 30% sucrose, and sectioned frozen on a sliding microtome at 40 µm or 60 µm. All primary and secondary antibodies were diluted in PBS containing 0.25% Triton X-100, 10% horse serum, and 2% BSA. The following antibodies were used: goat anti-mCherry (1:500, Antibodies Online #ABIN1440058), chicken anti-GFP (1:1000, Aves Labs #GFP-1020), rabbit anti-parvalbumin (1:5000, Swant #PV-27), guinea pig anti-parvalbumin (1:1,000, Synaptic Systems #195-004), Wisteria Floribunda Agglutinin (1:200, Sigma Aldrich #L-1516), rabbit anti-somatostatin (1:3,000, BMA Biomedicals #T-4103), mouse anti-PGC-1⍺ (1:200, Santa Cruz #sc-518025), mouse anti-Kv1.1 (1:1,000, Antibodies Incorporated #75-105), mouse anti-Kv3.1 (1:1,000, Antibodies Incorporated #75-041), mouse anti-ERRγ (1:100, R&D Systems #PP-H6812-00), rabbit anti-MEF2c (1:100, ProteinTech #18290-1-AP). We used Alexa Fluor-conjugated (1:400, ThermoFisher), donkey-anti-guinea pig 647 (1:250, Jackson ImmunoResearch #706-605-148), donkey-anti-chicken 488 (1:200, Jackson ImmunoResearch #703-545-155), donkey anti-rabbit 405 (1:200, Abcam #ab175651), Streptavidin 555 (1:400, ThermoFisher #S21381) and Streptavidin 647 (1:400, Jackson ImmunoResearch #016-600-084) secondary antibodies. Sections were finally counterstained with DAPI (5 µM) and mounted with Mowiol-Dabco mounting medium (Sigma).

#### Single-molecule fluorescent in situ hybridization

All single-molecule fluorescent *in situ* hybridization solutions were prepared in RNAse-free water and PBS. Mice were perfused as described above, and brains were postfixed overnight at 4°C, cryoprotected in 15% sucrose followed by 30% sucrose, and sectioned frozen on a sliding microtome at 30 µm. Sections were mounted on RNAse-free SuperFrost Plus slides (ThermoFisher) and probed against target RNAs using the RNAscope Multiplex Fluorescent Assay v2 according to the manufacturer’s protocol (ACDbio #323110). The following probes were used: Mm-*Lhx6*-C2 (#422791-C2), Mm-*Lhx6*-C3 (#422791-C3), Mm-*Sst*-C3 (#404631-C3), Mm-*Ppargc1a*-C1 (#402121), Mm-*Bcan*-O2-C2 (#531761-C2), Mm-*Cox6a2*-C3 (#461371-C3), Mm-*Esrra*-O2-C2 (#534441-C2), Mm-*Esrrg*-C3 (#495121-C3), Mm-*Reln*-C2 (#405981-C2), Mm-*Tac1*-C3 (#410351-C3), Mm-*Slc17a6-C2* (#319171-C2), Mm-*Slc17a7-C1* (#416631). When required, the RNAscope protocol was then followed by immunohistochemistry against mCherry (goat anti-mCherry, 1:250, Antibodies Online) to stain cells infected with adeno-associated viruses expressing mCherry or against GFP (using a combination of two rabbit anti-GFP antibodies, 1:500, Aves Labs #GFP-1020 and Abcam #ab13970) to reveal the endogenous GFP signal of MGE-derived cells labeled with an RCE reporter. Mounted sections were washed with PBS and blocked for 30 min in PBS containing 10% horse serum and 2% BSA. The sections were then incubated overnight at 4°C with the primary antibody. The next day, the sections were washed three times in PBS and incubated with a secondary antibody for 2 h at room temperature. Mounted sections were finally counterstained with DAPI (5 µM) before drying and applying Mowiol-Dabco mounting medium (Sigma). Primary and secondary antibodies were diluted in PBS with 10% horse serum and 2% BSA.

### Single-cell RNA sequencing

#### Preparation of single-cell suspension

*Lhx6-Cre;Ppargc1a^+/+^*and *Lhx6-Cre;Ppargc1a^F/F^* animals, injected in both hemispheres with *AAV8-hSyn-dio-tdTomato* viruses at P1, were processed separately in three control and three mutant batches. For each animal, the somatosensory cortex of both hemispheres was dissociated into a single-cell suspension. Mice were deeply anesthetized with sodium pentobarbital by intraperitoneal injection and transcardially perfused with ice-cold oxygenated cutting solution (87mM NaCl, 2.5mM KCl, 1.25mM NaH_2_PO_4_, 26 mM NaHCO_3_, 75mM sucrose, 20mM D-glucose, 0.5mM CaCl_2_*2H_2_O, 4mM MgSO_4_*7H_2_O). The brain was extracted and incubated for three minutes in an ice-cold oxygenated cutting solution. The somatosensory cortex was then dissected into chunks in icy oxygenated cutting solution, and the tissue was dissociated using the Papain Dissociation System (Worthington #LK003160). After 30 minutes of incubation at 37°C with papain solution, the tissue chunks were washed with an oxygenated cutting solution and were manually and slowly dissociated using a 10mL pipette and a 1mL pipette tip. All solutions used during this process were oxygenated for at least 10 minutes with 95% O_2_ and 5% CO_2_. After papain incubation, all steps were performed on ice and using low retention filter tips (VWR). Oxygenation and a short time of dissection are crucial for the viability of the cells in suspension. After dissociation, small tissue chunks may remain, and these were discarded by filtering through a 100µm cell strainer (Fisher Scientific #11587522). The debris were removed by overlaying the single-cell suspension on top of a 10% Percoll solution (Sigma #P4937, diluted in oxygenated cutting solution and 1.5M NaCl). After a 10-min 500g centrifugation at 4°C, the pelleted single-cells were resuspended in cutting solution with 2% BSA, filtered through a 70µm cell strainer (Fisher Scientific #11597522), and transferred into a FACS tube (Falcon #352063) precoated with 3% BSA overnight.

#### Fluorescence-activated cell sorting

Samples were sorted at 4°C on BD FACS Aria II or Aria III Cell sorters for no longer than 40 min and using a 130 µm nozzle to increase cell viability. DAPI was added to the single-cell suspension to discard dead cells. Viable tdTomato+ single cells were collected into 1 mL of cutting solution with 2% BSA in LoBind tubes (SLS #Z666505) precoated with 3% BSA overnight. Collection tubes were centrifugated for 5 min at 4°C at 500 g, and the supernatant was discarded until 45 µL were left in the tube. Cells were resuspended, and cell number and viability were assessed using Trypan Blue and a hemacytometer for cell counts. We only carried out single-cell RNA-sequencing in experiments with viability above 60%. Between 3,000-5,000 cells were loaded for each sample onto a 10X Genomic single-cell chip/machine for single-cell capture and cDNA library preparation. RNA-seq was performed in an Illumina NextSeq 2000 platform.

### Bioinformatic analyses of single-cell RNA sequencing

#### Data pre-processing

Illumina BCL output files were demultiplexed into FASTQs and used as inputs to the Cell Ranger count pipeline (v7.0.1). Alignment was performed to the mm10 genome and intronic reads excluded, but the pipeline was otherwise run with default parameters. The filtered feature barcode matrix was used as input for further analysis in Seurat (v4.3.0) in Rstudio (v2023.06.2+561).

#### Quality control

Upon initial creation of the Seurat object, genes detected in fewer than 3 cells and cells with fewer than 200 reads were discarded. Mitochondrial, ribosomal, and sex-specific genes were not excluded. In subsequent quality control, cells were retained if they had <10% ribosomal reads. Lower and upper limits for unique gene counts and lower limits for the percentage of mitochondrial reads were established for each sample individually (Figure S3). The total number of cells retained after filtering were 1489, 2797, and 982 cells for control replicates and 772, 991, and 1376 cells for mutant replicates.

#### Normalization, scaling, and variable feature detection

Cell cycle scores were calculated using the *CellCycleScoring* function. Sample read counts were subsequently normalized and variance-stabilized using *SCTransform*, regressing out percentages of mitochondrial and ribosomal reads and cell cycle score difference (calculated by subtracting the G2M score from the S score). This function was also used to identify the top 3000 variable features.

#### Data integration

Replicates within each condition were merged, and the pooled control and mutant datasets were individually subjected to *SCTransform* normalization. The same variables were regressed as above. The control and mutant datasets were then integrated according to the *SCTransform* integration workflow (*SelectIntegrationFeatures* with the top 3,000 variable features, *PrepSCTIntegration*, followed by *FindIntegrationAnchors* and *IntegrateData* specifying SCT as the normalization method).

#### Dimension reduction and clustering

Dimension reduction and clustering were carried out using the standard Seurat workflow (*RunPCA* with all variable features, *RunUMAP* and *FindNeighbors* with all 50 dimensions, and *FindClusters* at 1.5 resolution). Markers for each cluster were computed with the *FindAllMarkers* function. Only genes expressed in >10% of cells in at least one of the populations being compared were considered, and markers with adjusted *p* < 0.05 were taken to be differentially expressed. Of 27 clusters, 7 pertaining to non-neuronal or other undesired populations were excluded. The remaining clusters could be assigned PV+ or SST+ interneuron identity based on the expression of *Pvalb* and *Sst*, yielding a total of 1,376 and 995 control and mutant PV+ interneurons, and 3,098 and 2029 control and mutant SST+ interneurons, respectively. The control and mutant datasets were then split, *SCTransformed*, re-integrated according to the *SCTransform* integration workflow, and subjected to dimension reduction (*RunPCA* with all variable features, *RunUMAP* with 50 dimensions).

#### Differential gene expression analysis

DGE analysis was performed on RNA counts normalized according to default parameters, employing the *FindMarkers* or *FindAllMarkers* functions in Seurat. Unless specified otherwise, only genes expressed in >10% of cells in at least one of the populations being compared, and with average log_2_ fold change >|0.25| and adjusted p-value < 0.05, were considered. To find PV and SST-enriched genes at P21, we compared wild type PV+ and SST+ interneurons in our own dataset, using a log_2_ fold change threshold of >|0.35|. To find mature markers of different interneuron subclasses, we performed DGE analysis using a single-cell RNA-sequencing dataset from the adult mouse cortex^53^ and used a log_2_ fold change threshold of >|0.45|. To identify genes enriched in L4 and L5/6 PV+ interneurons, we used the Patch-seq dataset from the adult mouse cortex^70^ to infer the layer localization of different transcriptomic subtypes. Layer assignments were generalized to the entire Patch-seq dataset, but Chandelier cells (*Pvalb Vipr2*) and subtypes that could not be assigned to any layer (*Pvalb Calb1 Sst*, *Pvalb Gpr149 Islr*) were excluded from the analysis. PV+ interneuron subtypes assigned to L4 and L5/6 were used for head-to-head DGE analysis.

#### Assignment of layer identities

As described above, we used a Patch-seq dataset^70^ to infer layer localization of different PV+ interneuron subtypes. Read counts were subjected to *SCTransform* normalization as described for the P21 dataset. Layer identities were assigned from the reference dataset by label transfer (*FindTransferAnchors*, followed by *TransferData*), specifying 50 dimensions, *SCTransform* as the normalization method, and employing CCA dimension reduction under the assumption that all cortical layers are represented in our dataset.

#### Pathway enrichment analysis

Pathway enrichment analysis was performed with the *compareCluster* function from the *clusterProfiler* package (v4.4.2), using Reactome pathway terms.

### CUT&RUN

#### Preparation of nuclei suspension

The somatosensory cortex was dissected from P12 WT C57BL6 mice in ice-cold buffer HB (0.25 M sucrose, 25 mM KCl, 5 mM MgCl2, 20 mM Tricine-KOH pH 7.8, 1 mM DTT, 0.15 mM spermine, 0.5 mM spermidine). Cortices were homogenized by a Dounce homogenizer in the presence of 0.3% IGEPAL. The homogenate was then filtered through a 40 µm strainer (Fisher Scientific #11597522), centrifuged at 4°C for 5 min at 500 g, and the pellet was washed twice with 1% BSA solution. The pelleted nuclei were resuspended in CUT&RUN Wash buffer (20mM Tris-HCl pH 7.5, 150mM NaCl, 0.5mM spermidine, 0.1% BSA), and the shape of the nuclei was verified using Tryptan Blue and a hemacytometer. All steps were performed on ice and using LoBind pipette tips.

#### CUT&RUN

Nuclei were incubated with 2 μg of mouse anti-PGC-1⍺ antibody (Millipore #ST1202, two replicates) or 2 μg of mouse anti-ERRγ antibody (R&D Systems #PP-H6812-00, one replicate) in CUT&RUN Wash buffer for 10min with rotation at 4°C. 2 μg of rabbit anti-mouse IgG secondary antibodies (Abcam #ab6709) were then added to the nuclei solution, which was left to rotate for 10min at 4°C. Nuclei were then incubated without washing step with pA-MNase (1.6 µg/µl) at 4 °C with rotation for an additional 10 min. Nuclei were then washed and resuspended with CUT&RUN wash buffer. DNA-tethered cleavage was induced by adding CaCl_2_ at a final concentration of 2 mM on ice water for 10 min. The reaction was stopped by adding Stop buffer (200 mM NaCl, 20 mM EDTA, 4 mM EGTA, 50 µg/ml RNAse A (Thermo Scientific #EN0531), 40 µg/ml glycogen (Thermo Scientific #R0561)) at a 1-to-1 ratio. The shape of the nuclei was again verified using Tryptan Blue and a hemacytometer. To release the DNA fragments of interest, nuclei were incubated at 37 °C for 20 min. Solubilized chromatin of interest was separated from nuclei by centrifugation at 16,000 g at 4°C for 5 min and then digested with SDS (0.1% final concentration) and Proteinase K (0.2 mg/ml final concentration) at 70°C for 10 min. Released DNA fragments were then extracted with phenol:chloroform:isoamylalcohol (25:24:1) and 13,000 rpm centrifugation for 15 min at room temperature. The aqueous upper layer was transferred to a fresh tube, and NaCl was added at a final concentration of 250 mM. An equal volume of isopropanol was further added, and the DNA was left to precipitate for 2 h at -80°C. Precipitated DNA was then pelleted by centrifugation (13,000rpm for 15 min at 4°C), washed with 70% ice-cold EtOH, and resuspended in distilled water.

#### Library preparation

Libraries for sequencing were prepared using the NEBNext Ultra II library prep kit (New England Biolabs) following the protocol and thermocycling temperatures therein. Briefly, DNA ends from fragments harvested from the CUT&RUN experiment were repaired, 5’ ends phosphorylated and 3’ ends A tailed prior to ligation with NEBNext Adaptor, UNG was then used to linearize the ligated adapter. SPRIselect beads were used to deplete the unligated adaptor. Adaptor-ligated libraries were then barcoded by PCR following the manufacturer’s recommended protocol. Libraries were amplified for between (11-13) cycles and purified using SPRIselect beads prior to quantitation using Qubit and HS DNA tapestation. Individually barcoded libraries were then pooled and sequenced on an Illumina NextSeq 2000 instrument.

### Bioinformatic analyses of CUT&RUN

#### CUT&RUN data processing

Raw sequencing reads were trimmed using TrimGalore (v0.6.6) to remove adapters and low-quality reads. The trimmed fastq files were aligned to the mm10 genome using bowtie (v2.2.9) with the following parameters: --end-to-end --very-sensitive -- no-mixed --no-discordant --phred33 -I 10 -X 700. PCR duplicates were removed using Picard (v2.6.0), and bam and fragment bed files were generated using Samtools (v1.3.1) and Bedtools (v2.25.0), selecting fragment lengths under 1,000 bp. Peaks were called for the two replicates of mouse anti-PGC-1⍺ antibody (Millipore #ST1202) and the replicate of mouse anti-ERRγ antibody (R&D Systems #PP-H6812-00), using SEACR (Sparse Enrichment Analysis for CUT&RUN; v1.3), based on fragment bedgraph files generated with Bedtools *genomecov*. SEACR peaks were called as follows: SEACR_1.3.sh input_bedgraph 0.005 non stringent out_name. PGC-1⍺ peaks were retained if they were captured in both replicates. All peaks overlapping with mm10 blacklisted regions (ENCODE file ID: ENCFF547MET) were removed prior to further analysis. Transcription factor motifs were identified with HOMER (Hypergeometric Optimization of Motif EnRichment; v4.10) as follows: findMotifsGenome.pl input_bed mm10 out_dir -size given. CUT&RUN peaks on the mm10 genome were visualized using the IGV (v2.16.0) software.

#### Gene set enrichment analysis

To derive targets from CUT&RUN data, peaks were annotated to genes using the annotatePeak function from the ChIPseeker package (v1.32.0), using the TxDb.Mmusculus.UCSC.mm10.knownGene and org.Mm.eg.db databases and specifying promoter regions within 3000bp on either side of the transcription start site (TSS). Genomic targets of PGC-1⍺ and ERRγ were annotated as promoters, introns, exons, UTRs, or distal intergenic.

### Biochemistry

#### Nuclear fractionation

Cortices were rapidly dissected in ice-cold PBS, homogenized in cytosol lysis buffer (10 mM Hepes, 40 mM KCl, 3 mM MgCl_2_, 5% glycerol, 0.2% NP40, 1 mM DTT, cOmplete protease inhibitors, Sigma-Aldrich) and incubated for 30 min at 4°C with rotation. Following centrifugation at 500 g for 5 min at 4°C, the supernatant containing the cytosolic proteins was denatured at 80°C for 10 min in Laemmli sample buffer (80 mM Tris-HCl pH 6.8, 100 mM DTT, 8.7% glycerol, 2% SDS, 0.01% bromophenol blue). The pellet containing the nuclear proteins was washed in cytosol lysis buffer, resuspended in nuclear lysis buffer (20 mM Hepes, 420 mM NaCl, 1.5 mM MgCl_2_, 25% glycerol, 0.2 mM EDTA, 1 mM DTT, cOmplete protease inhibitors) and incubated for 30 min on ice with regular vortexing. Following centrifugation at 14,000 g for 10 min at 4°C, the supernatant containing the nuclear proteins was recovered. For Western blot, the nuclear fractions were denatured at 80°C for 10 min in Laemmli sample buffer. Protein concentrations were measured using Pierce 660 nm Protein Assay (ThermoFisher).

#### Co-immunoprecipitation

Nuclear fractions from P12 C57BL/6 mice were diluted in co-immunoprecipitation buffer (20 mM Hepes, 90 mM NaCl, 1 mM MgCl_2_, cOmplete protease inhibitors) and subsequently incubated overnight at 4°C with 8 µg of either rabbit anti-Mef2c (Proteintech #18290-1-AP) or rabbit IgG isotype control (Abcam #ab27478). 25 µl of Dynabeads Protein G slurry (Invitrogen) was washed in co-immunoprecipitation buffer and added to the samples for 3 h at 4°C with gentle rotation. Beads were washed six times in wash buffer (20 mM Hepes, 150 mM NaCl, 1% NP40, 1% glycerol, 1 mM MgCl_2_, cOmplete protease inhibitors), resuspended in Laemmli sample buffer, and samples were denatured at 80°C for 10 min.

#### Western blotting

20 µg of denatured protein extracts (or the whole denatured co-immunoprecipitation samples) were separated by SDS-PAGE using 10% acrylamide gels for 2 h at 120V and transferred onto methanol-activated PVDF membranes at 300 mA for 2 h on ice. Membranes were blocked with 5% non-fat milk in TBS-T (20 mM Tris-HCl pH 7.5, 150 mM NaCl, and 0.1% Tween20) for 1h at room temperature and then probed with primary antibodies in blocking buffer overnight at 4°C. The next day, membranes were washed in TBS-T and incubated with HRP-conjugated secondary antibodies for 1 h at room temperature. Depending on the amount of protein of interest in the samples, membranes were incubated with either Immobilon Western Chemiluminescent HRP substrate (Millipore), SuperSignal West Pico, or West Femto Chemiluminescent Substrates (ThermoFisher). Protein levels were visualized by chemiluminescence with an Odyssey FC (Li-Cor) and quantified with Image Studio Lite. For quantification, the densitometry of the band of interest was normalized to that of actin. The following primary antibodies were used: mouse anti-PGC-1⍺ (1:200, Santa Cruz #sc-518025), rabbit anti-MEF2c (1:500, Proteintech #18290-1-AP), mouse anti-actin-peroxidase (1:20000, Sigma-Aldrich #A3854). The following HRP-conjugated secondary antibodies were used: anti-mouse-peroxidase (1:5000, Invitrogen #31444), anti-mouse-peroxidase (1:1000, Abcam #ab131368) and anti-rabbit-peroxidase (1:1000, Abcam #ab131366).

### Luciferase assay experiments

We constructed luciferase reporter vectors by cloning promoter regions of the *Bcan*, *Cplx1*, and *Kcan1* genes in a pGL4.11 plasmid (Promega) by Gibson assembly (New England Biolabs) using the following primers: *Cplx1* Forward: CCAGAACATTTCTCTGGCCTCTTACTCCCCACCCCTAGC; *Cplx1* Reverse: ACAGTACCGGATTGCCAAGCGCCGGTTACCTTCC. *Bcan promoter 1* Forward: GGCCTAACTGGCCGGTACCCAGAGCGCTGCTTAGCAGG*; Bcan promoter 1* Reverse: CCAACAGTACCGGATTGCCATAGAGACGGGTTCCTGTGTATGAAAGAC. *Bcan promoter 2* Forward: GGCCTAACTGGCCGGTACGTGTTGGCACCATCTTACATGTGG; *Bcan promoter 2* Reverse: CCAACAGTACCGGATTGCCACTTGCGAGGTCCCCTCC; *Kcna1* Forward: GGCCTAACTGGCCGGTACAGCCACTTGGGAGGATCC; *Kcna1* Reverse: CCAACAGTACCGGATTGCCACCTCAGTCCTGCCTTCCGA. HEK293T (embryonic kidney) cell lines, obtained from the American Type Culture cells, were maintained with DMEM medium supplemented with L-glutamine, 10% fetal bovine serum (Sigma-Aldrich) and 1% penicillin-streptomycin (Invitrogen). Cells were allowed to grow at 37 °C with 5% CO_2_. HEK293T cells were seeded in 12 wells plates and transfected (Fugene HD, Promega) the next day with the luciferase reporter vectors (100 ng), the expression vectors (100 ng each), and the pRL-TK plasmid to express *Renilla* as an internal control (2 ng). After 16 hours, cells were washed with PBS, and luciferase assays were performed in triplicates using the Rapid Detection Firefly Luciferase Activity assay system (Promega), following the manufacturer’s instructions, with a GloMax plate reader (Promega). Firefly luciferase activities were normalized based on the *Renilla* luciferase activity in each well.

### Histological image acquisition and analysis

Images were acquired with an SP8 confocal microscope (Leica TCS SP8). Samples from the same experiment were imaged and analyzed in parallel, using the same laser power, photomultiplier gain, and detection filter settings.

#### Cell densities and proportions

Imaging was performed at 12-bit depth using a 10X objective, except for PGC-1⍺ and ERRγ immunostainings where images were acquired with a 20X objective. Images following immunohistochemistry experiments were acquired with a 1024×1024-pixel resolution, while imaging of *in situ* hybridization experiments was performed with a 2048×2048-pixel resolution. Parameters of 0.75 digital zoom and 400 Hz acquisition speed were used. Between 7 and 10 sections per brain were imaged and analyzed after *in situ* hybridization or immunohistochemistry. Manual cell counting was then performed across layers of the somatosensory cortex using the FIJI (Image J) software.

#### Intensities of PV and WFA and PGC-1⍺

Imaging was performed at 16-bit depth using a 63X objective with water immersion. Images were acquired with a 2048×2048-pixel resolution, 1 or 1.5 digital zoom, and 600 Hz acquisition speed. Z-stacks were taken with a Z-step size of 1 (except for measuring PGC-1⍺ levels in P10 and P12 MGE-derived cells, where a Z-step size of 2 was used) to consider the volume of a selected cell to image and analyze. A sum Z-projection was then performed for each image on FIJI.

In the excitatory DREADDs (hM3Dq-mCherry) experiments, putative PV+ cells in L2/3 of the somatosensory cortex labeled with mCherry or GFP were imaged, maintaining a consistent depth to the pial surface. In the absence of DAPI, L2/3 was recognized from L4 by the absence of dense PV+ processes. In the inhibitory DREADDs (hM4Di-mCherry) experiments, putative PV+ cells of L5 of the somatosensory cortex labeled with mCherry or GFP were imaged. Similarly, in the absence of DAPI, L5 was recognized from L4 and L6 by the global pattern of PV expression. A minimum of 15 DREADDs-infected mCherry+ cells and 15 DREADDs-uninfected GFP+ cells were analyzed per brain in Matlab, using a custom script. Briefly, for each image, background subtraction was applied. The contour of an mCherry+ prospective PV+ cell (identified by the lack of SST expression) was then used to draw a ROI. The mean pixel value of this ROI was measured to provide the PV intensity level of the DREADD-infected cells. PV intensity levels of DREADDs-uninfected mCherry-GFP+ prospective PV+ cells (identified by the lack of SST expression) were also measured by drawing a ROI following the contour of the GFP signal. The PV intensity levels of the same number of DREADDs-infected and -uninfected prospective PV+ cells were measured per image. We calculated the median value of PV intensity levels for DREADDs-infected and -uninfected prospective PV+ cells separately per brain. To account for the different speeds of PV+ interneuron maturation between animals, PV intensity levels of DREADDs-infected prospective PV+ cells were normalized for each animal with the PV intensity levels of DREADDs-uninfected prospective PV+ cells in the same brain. These images were also used to count the number of SST+ cells infected by DREADDs expressing viruses (SST+ mCherry + cells) and calculate the rate of infection of this virus in MGE-derived cells. WFA intensity levels were measured following the same methodology.

In experiments assessing the role of excitatory innervation on the maturation of PV+ interneurons using *ErbB4^F/F^;Ai9* mice and *ErbB4^+/+^;Ai9* mice injected with *E2-Cre* AAV8 viruses, recombined tdT+ PV+ cells in L5/6 of the somatosensory cortex were imaged. Because the expression of Cre leaks to excitatory neurons and perhaps other interneurons, we only considered PV+ cells that were expressing PV. We analyzed a minimum of 15 recombined tdT+ PV+ cells per brain, as well as a minimum of 15 unrecombined PV+ cells present in each image, using a custom Matlab script. Briefly, for each image, background subtraction was applied. The contour of PV+ cells present in the image was drawn based on the PV signal, delineating a ROI. For each image, we identified recombined tdT+ PV+ cells and unrecombined tdT-PV+ cells. The mean pixel value of the ROI was measured to provide the PV intensity level of each PV+ cell (tdT+ and tdT-). We calculated the median value of PV intensity levels for tdT+ and tdT-PV+ cells separately, per image and then per brain. This value for tdT+ PV+ cells was normalized by the value for tdT-PV+ cells per brain to account for the differences in PV interneuron maturation speed between brains. PGC-1⍺ intensity levels were measured using the same methodology; only intensity levels in the nucleus (delineated using the DAPI signal) and the cytoplasm of each cell were measured.

In the experiments measuring PGC-1⍺ intensity levels in prospective PV+ interneurons at P10 and P12 in *Lhx6-Cre;RCE* mice, these interneurons were recognized by the expression of the GFP reporter and the lack of SST expression. Putative PV+ interneurons of L2/3, L4, and L5 of the somatosensory cortex (20 cells per layer) were imaged and analyzed using a custom-made Matlab script. Briefly, for each image, background subtraction was applied. The somatic contour of a prospective PV+ cell was drawn using the GFP signal to delineate a ROI. The mean pixel value of this ROI was measured to provide the PGC-1⍺ intensity level of this cell. Intensity levels in the nucleus (delineated using the DAPI signal) and the cytoplasm of each cell were also measured.

#### RNAscope dot count

For *Esrra* and *Esrrg* mRNA particle analysis, imaging was performed at 16-bit depth using a 63X objective with water immersion. Images were acquired with a 2048×2048 pixel resolution, 1 digital zoom, and 600 Hz acquisition speed. Five S1 L2/3 and five S1 L4 ROIs were imaged per brain. *Esrra* and *Esrrg* levels were measured in a minimum of 30 MGE-derived cells per brain. In this experiment, MGE-derived cells were identified by the expression of tdT. For all these experiments, the number of *Esrra* and *Esrrg* mRNA particles was determined using a custom Matlab script. Briefly, for each image, background subtraction was applied. The contour of tdT+ cells was then used to draw a ROI. Within each ROI, the area of total *Esrra* and *Esrrg* signals was measured. This area was divided by the area of a single mRNA particle (∼0.16 µm^2^), which estimates the number of *Esrra* and *Esrrg* particles present in the ROI of interest.

#### Mito-EmGFP counts

Imaging was performed at 16-bit depth using a 63X objective with water immersion. Single-plane images were acquired with a 2048×2048 pixel resolution, 1.25 digital zoom, and 600 Hz acquisition speed. Twelve ROI in L2/3 and nine ROI in L4 of the somatosensory cortex were imaged per brain. Mito-EmGFP counts were measured in a minimum of 60 MGE-derived cells per brain (30 SST+ and 30 prospective PV+ interneurons, the latter identified by the lack of SST expression, using a custom Matlab script. Briefly, for each image, general background subtraction was applied. The contour of infected cells was drawn using the mito-EmGFP signal to delineate the ROI around the soma of each cell. Similarly, we drew the contours of the nucleus of each cell using the mito-EmGFP signal. We measured the levels of GFP intensity in the nucleus of each cell, which accounts for background GFP intensity levels, whose value depends on the efficacy of infection of the mito-EmGFP expressing virus per cell (more infection, more mito-EmGFP expression, more residual GFP present in the nucleus and cytoplasm). We then subtracted this background intensity value from the somatic ROI per cell. We also removed the nucleus from the somatic ROI to consider from now on only a cytoplasmic ROI. Within each cytoplasmic ROI, the area occupied by the mito-EmGFP signal was measured, and the proportion of the cell cytoplasmic area occupied by the mito-EmGFP signal was calculated.

#### Synapses

Imaging was performed at 8-bit depth using a 100X objective with oil immersion. Single-plane images were acquired with a 2048×2048 pixel resolution, 2.2 digital zoom, and 600 Hz acquisition speed. We compared the density of Homer+ Bassoon+ excitatory synapses contacting the soma of tdT+ infected and neighboring tdT-uninfected PV+ cells in L5/6 of *ErbB4^F/F^;Ai9* mice. Only PV+ cells with detectable levels of PV expression were recognized and used for quantification. Between 15 and 30 L5/6 PV+ cells were analyzed per brain. We used a custom macro in FIJI to measure Homer-Bassoon synaptic density. Background subtraction, Gaussian blurring, smoothing, and contrast enhancement were first applied in all channels. Cell somas were drawn manually based on intensity levels of PV staining to create a mask of the soma surface and measure its perimeter. Presynaptic and postsynaptic boutons were detected automatically based on thresholds of intensity. The threshold for Homer and Bassoon were selected from a set of random images before quantification, and the same thresholds were applied to all images from the same experiment. The “Analyze Particles” and “Watershed” tools were applied to the Homer and Bassoon channels, and a mask was generated with a minimum particle size of 0.06 to detect colocalized Homer and Bassoon puncta. These puncta were defined as synaptic boutons when they were located up to 0.3μm^2^ inside or 0.4μm^2^ outside the soma perimeter.

### Statistical analyses

All statistical analyses were performed using Prism 9 (GraphPad Software). No statistical methods were used to predetermine sample sizes. Sample sizes were chosen based on previous publications in the field. Biological replicates (n values are different samples derived from different brains from different litters) were analyzed to assess the biological variability and reproducibility of data. Experimental mice from all genotypes or conditions were processed together. Samples were tested for normality using the Shapiro–Wilk normality test. Unless otherwise stated, data were analyzed by t-test or ANOVA followed by post hoc Tukey’s analysis for comparisons of multiple samples. Differences were considered significant when p values < 0.05. Data are presented as mean ± s.e.m. Statistical details of experiments are described in figure legends.

